# Genome-wide identification and characterization of the HSP gene superfamily in apple snails (Gastropoda: Ampullariidae) and expression analysis under temperature stress

**DOI:** 10.1101/2022.06.30.498069

**Authors:** Yue Gao, Jia-Nan Li, Jia-Jia Pu, Ke-Xin Tao, Xing-Xing Zhao, Qian-Qian Yang

## Abstract

Heat shock proteins (HSPs) play important roles in the response to various stresses as molecular chaperones. Apple snails from the family Ampullariidae have become economically important due to several species mainly from the genus *Pomacea* were invasive. The recent release of the genomes of ampullariids (*P. canaliculata, P. maculata, Lanistes nyassanus*, and *Marisa cornuarietis*) has opened the opportunity for a comprehensive analysis of the HSP superfamily. We identified that the number of HSP from *P. canaliculata* (PcaHSPs) was greater than that from the other three species. A total of 42 PcaHSPs were distributed on 12 chromosomes and were classified into the families of HSP90, HSP70, HSP60, HSP40, HSP20, and HSP10. Each family formed a monophyletic clade on the phylogenetic trees, except for the HSP40 family. We identified tandem duplication of paralogous genes in PcaHSP70 and PcaHSP20. The RNA-seq data show that the expression profiles of PcaHSPs in different tissues have similar patterns, except that several PcaHSP20 genes revealed tissue-specific expression levels. Moreover, we identified that there were more HSP genes with stronger induction levels in response to hot than cold stress. Our findings will be helpful for future studies on stress response and adaptation focusing on HSPs in apple snails.

## 1. Introduction

Heat shock proteins (HSPs) are a family of conserved proteins that can be rapidly synthesized and help regulate the response to protect organisms from cellular damage induced by environmental stress, such as temperature, drought and heavy metals [1-3]. HSPs can be generally divided into seven families based on their gene structures and protein motifs, including HSP100, HSP90, HSP70, HSP60, HSP40, HSP10, and sHSPs (small HSPs), according to their molecular weights [4, 5]. Hsp100 is not found in animals but in bacteria, yeast, and plants [6]. sHSPs are of small molecular mass in the range of 12-42 kDa with a conserved α-crystallin domain (or HSP20 domain) of 90-100 amino acids in length [7, 8]. HSP70 is one of the most conserved protein families and can be classified into induced heat shock protein (HSP70) and heat shock cognate protein 70 (HSC70) based on expression profiles [2, 9]. HSP genes from each family have been extensively studied in various taxa to reveal their expression patterns for tolerance against environmental stresses. In particular, HSP90 and HSP70 have received the most research attention in gastropods, such as the apple snail *Pomacea canaliculata* [10], the blood fluke planorb *Biomphalaria glabrata* [11], and the variously colored abalone *Haliotis diversicolor* [12].

Apple snails are freshwater species in the family Ampullariidae (Molluska: Caenogastropoda), which has approximately 120 species in nine existing genera with diverse geographical distribution patterns [13]. The genera *Pomacea, Marisa, Felipponea*, and *Asolene* are native to the New World, whereas *Pila, Lanistes, Afropomus, Saulea*, and *Forbesopomus* are native to the Old World [13]. The asymptotic biological and physiological traits approaching amphibiousness, such as the diversity in respiration morphology and desiccation resistance, oviposition location and egg morphology, have facilitated Ampullariidae as a good molluskan model for understanding the evolutionary transition from aquatic to terrestrial [14, 15]. Moreover, apple snails have become economically important due to several *Pomacea* species; notably, *P. canaliculata* and *P. maculata* have been introduced out of their native ranges and have become highly invasive pests in aquatic ecosystems and agriculture [16]. Characterizing the key gene families related to responses to environmental stresses, such as HSPs, will provide basic knowledge for understanding the underlying process of adaptation in apple snails and thus offer insights into their evolution and invasion.

For example, the environmental temperature plays a crucial role in the survival [17], growth [18], development [19], and reproduction [20] of apple snails, which therefore affects their geographic distribution [19, 21]. The optimum temperature of *P. canaliculata* is 25°C, which maximizes the growth and survival rates [18]. Below or above the optimum temperature ranges can affect their biology, ethology, and physiology to different degrees [22, 23]. For example, the oviposition of *P. canaliculata* stops when the temperature is <15° or >35°, while its activity may stop and gradually start dormancy when below 10° or above 39° [24, 25].

Five HSP genes have been characterized from *P. canaliculata*, including two HSP70 genes [26, 27] and one gene from each HSP90 [10], HSP60 [10], and HSP40 [28] family. In recent years, the report of high-quality, chromosome-level assembly genomes has allowed the identification and better understanding of HSPs in a couple of species, such as the sea squirt *Ciona savignyi* [29], the mosquito *Anopheles sinensis* [30], and the whitefly *Bemisia tabaci* [31]. The recent release of the whole genome sequences of four apple snail species, *P. canaliculata, P. maculata, Marisa cornuarietis*, and *Lanistes nyassanus* [15], has opened the opportunity for a full view of the HSPs. In this study, we aimed to characterize the HSP genes of apple snail species from genome-wide analysis and to reveal the expression profile of HSP genes in different tissues from RNA-seq data of *P. canaliculata*. Moreover, we will investigate the expression patterns of HSP genes in *P. canaliculata* under heat and cold stress to better interpret their roles in temperature tolerance.

## 2. Materials and methods

### 2.1 Identification of HSP genes in apple snails

The published genomes of four apple snail species, namely, *P. canaliculata, P. maculata, Lanistes nyassanus*, and *Marisa cornuarietis* were used to identify the HSPs [15]. The genome data and annotation files were accessed from https://datadryad.org/stash/dataset/doi:10.5061/dryad.15nd8v3. We retrieved all 130 available HSP protein sequences of gastropods from the National Center for Biotechnology Information (NCBI) protein database up to 2021-02-22 to form a query profile, which included 30 HSPs of the California sea hare *Aplysia californica*, 40 HSPs of the blood fluke planorb *Biomphalaria glabrata*, 31 HSPs of the owl limpet *Lottia gigantea*, 4 HSPs of the Japanese disc abalone *Haliotis discus hannai*, and 24 HSPs of *P. canaliculata* (Appendix S2: Table S1).

HSPs of *P. canaliculata* (PcaHSPs), *P. maculata* (PmaHSPs), *L. nyassanus* (LnyHSPs), and *M. cornuarietis* (McoHSPs) were obtained based on the multiple sequence alignment of the query profile in the genomic data carried out (-db all - outfmt 7 -out blast_out.tab -evalue 1e-20) by the Blastp program. Then, we searched the obtained HSPs on the ir-HSP [32] online server to verify the sequences and their family classification. The Pfam protein domain database (https://pfam.xfam.org/) was used to confirm the identified sequences containing functional conserved domains related to each HSP family.

The chromosomal localization of PcaHSPs was analyzed using the MG2C tool (http://mg2c.iask.in/mg2c_v2.0) according to the physical map described in the genome annotation files (.gff) of *P. canaliculata* [33]. We analyzed the duplication events of the PcaHSPs using the multiple collinear scanning toolkit (MCScanX) [34].

### 2.2 Phylogenetic analysis and classification of HSP genes

We performed multiple sequence alignment for the identified HSPs using the Muscle algorithm by MEGAX [35] and reconstructed the phylogenetic trees based on both the neighbor joining (NJ) and maximum likelihood (ML) methods. The NJ trees were based on the p-distance model and 1,000 bootstrap replications using MEGAX [35]. The ML trees were reconstructed using IQ-Tree 1.6.12 (-m TEST -bb 1000 -nt AUTO) [36].

### 2.3 Characteristics of PcanHSPs

We computed the amino acid length (AA), theoretical isoelectric point (pI), and molecular weight (Mw) using online ProtParam tools on the ExPASy website (https://www.expasy.org/protparam/) and predicted the subcellular localizations using the WoLF PSORT online server (http://psort.hgc.jp/).

To analyze the gene structures of PcanHSPs, we used the Gene Structure Display Server (GSDS) (http://gsds.cbi.pku.edu.cn/) to analyze the structure of the PcanHSP genes and display the numbers and positions of coding sequences (cds) and introns according to the genome annotation files (.gff) of *P. canaliculata*.

The conserved motifs of the PcanHSP proteins were predicted using the MEME online web server (http://meme-suite.org/) with the following parameters: zero or one per sequence, a maximum number of motifs of 20, and an optimum motif width from 30 to 100 sites [37].

### 2.4 Expression profiles of PcaHSPs in different tissues

We analyzed the expression profiles of the PcaHSPs based on the published RNA-seq data from tissues of hepatopancreas, kidney, gill, ovary, and testis[38], which was under the GenBank accession numbers SRR6429132, 133, 140-146, 153, 154, and 159-162 (Appendix S2: Table S2). The RNA-seq data were normalized using the scale of fragments per kilobase of transcript, per million mapped reads (FPKM) to reflect that a single sequenced molecule can generate two reads but came from a single cDNA fragment [39]. Then, the digital expression profile of PcaHSPs in the form of a heatmap was constructed using TBtools based on the transformed data of the log10 (FPKM+1) values.

### 2.5 Gene expression patterns of PcaHSPs under temperature stress

The snail population of *P. canaliculata* used for the experiment has been kept in our lab (25±1 °C, 14 L: 10D) for two years since collection from West Lake, Zhejiang Province of China (120°14′02″E, 30°25′70″N). We selected healthy females with a shell height of 15±2 mm and exposed them to hot (36°) and cold (9°) stress for 2 h and then returned them to normal temperature (25°) for 1 h to accumulate the expression of HSPs as previously described [10]. The control group of snails was kept at 25°C for 3 h. Hepatopancreas tissues from individuals were stored in liquid nitrogen immediately after dissection and transferred to -80 °C for preservation.

We extracted total RNA using an RNAprep Pure Tissue Kit (Tiangen, China) following the manufacturer’s protocol and then used DNase I to remove the remaining genomic DNA. We used the PrimeScript™ RT Reagent Kit with gDNA Eraser (TaKaRa, Japan) to synthesize first-strand cDNA from 2.5 μg of total RNA. The reaction conditions were 37°C for 2 min, 85°C for 5 s, and 4 °C for 5 min.

We designed the primers for PcaHSPs by Primer-BLAST (https://www.ncbi.nlm.nih.gov/tools/primer-blast/) and synthesized them at Shanghai Sunny Biotechnology Co., Ltd. (Shanghai, China) (Appendix S2: Table S3). After verifying the specificity of these primers with each sample from the three temperature treatments, we performed qPCR using the StepOnePlus Real-Time PCR System (BIO-RAD, USA) using SYBR@ Premix Ex TaqTM (TaKaRa, Japan) with *β-actin* as the internal control gene [10]. The conditions were 95°C for 30 s followed by 40 cycles of 95°C for 5 s and 53-62°C for 34 s (Appendix: Table S3).

The relative expression levels of the PcHSPs were calculated using the 2^−^ △△Ct method. We performed statistical analysis using data from three separate cDNA sets from three separate biological samples. Then, Mev 4.9.0 software [40] was used to generate the expression profiles.

## 3. Results

### 3.1 Identification and classification of HSPs in apple snail species

We identified a total of 42 potential HSP orthologs from the genomes of *P. canaliculata*, which was the highest number among the four apple snail species. There were 36, 32, and 40 HSP orthologs identified from the genomes of *P. maculata, M. cornuarietis*, and *L. nyassanus*, respectively (Table 1, Appendix S1). All the identified HSP orthologs were confirmed to be HSP proteins by the ir-HSP server and Pfam database. The HSP genes in the four apple snails were classified into the families of HSP90, HSP70, HSP60, HSP40, HSP20 and HSP10 based on the Pfam database. There were abundant members in the HSP70, HSP40, and HSP20 families, two or three members in HSP90, but only one member in each of the HSP60 and HSP10 families in all species (Table 1).

**Table 1.**
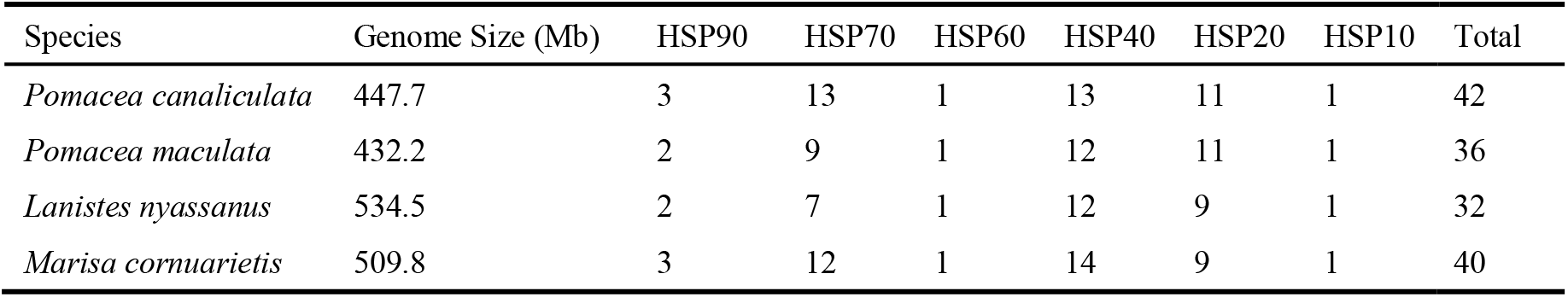
HSP genes identified in the genomes of four apple snail species

| Species | Genome Size (Mb) | HSP90 | HSP70 | HSP60 | HSP40 | HSP20 | HSP10 | Total |
| --- | --- | --- | --- | --- | --- | --- | --- | --- |
| *Pomacea canaliculata* | 447.7 | 3 | 13 | 1 | 13 | 11 | 1 | 42 |
| *Pomacea maculata* | 432.2 | 2 | 9 | 1 | 12 | 11 | 1 | 36 |
| *Lanistes nyassanus* | 534.5 | 2 | 7 | 1 | 12 | 9 | 1 | 32 |
| *Marisa cornuarietis* | 509.8 | 3 | 12 | 1 | 14 | 9 | 1 | 40 |

### 3.2 Genomic location and characteristics of PcaHSPs

We renamed the PcaHSPs in each family with continuous numbers (e.g., PcaHSP90-1) according to their orders of locations on the chromosomes (Fig. 1). Thirty-eight PcaHSPs were located on 12 of the 14 chromosomes of *P. canaliculata* (i.e., Chr03–Chr14), while the other four (PcaHSP90-3, PcaHSP70-13, PcaHSP40-13, and PcaHSP20-11) were identified on the scaffolds that were not assembled to chromosomes (Table 2; Fig. 1). Chr03, Chr07, and Chr10 have seven to nine PcaHSP genes. Chr03 has the most abundant distribution, with two PcaHSP90, one PcaHSP70, three PcaHSP40, and three PcaHSP20 genes. There were two and three PcaHSPs on Chr13 and Chr04, respectively, and only one PcaHSP on each of the other chromosomes.

**Table 2.** The chromosomal location, amino acids, molecular weight, isoelectric point and subcellular localization of HSPs in *P. canaliculata* (PcaHSP). AA, amino acids; MW, molecular weight; pI, isoelectric point

| Family | Gene Name | Gene ID | chr/<br>contig | AA | MW<br>(kDa) | pI | Subcellular localization |
| --- | --- | --- | --- | --- | --- | --- | --- |
| HSP90 | PcaHSP90-1 | Pca_57_9.24 | Chr03 | 726 | 83.59 | 4.96 | cytoplasm |
|  | PcaHSP90-2 | Pca_148_2.81 | Chr03 | 352 | 40.38 | 6.12 | cytoplasm |
|  | PcaHSP90-3 | Pca_1461_0.3 | Pca_1461 | 506 | 58.41 | 4.99 | nucleus |
| HSP70 | PcaHSP70-1 | Pca_2_20.24 | Chr03 | 850 | 95.90 | 5.14 | cytoplasm |
|  | PcaHSP70-2 | Pca_60_13.29 | Chr04 | 656 | 71.52 | 7.04 | mitochondrion |
|  | PcaHSP70-3 | Pca_15_37.41 | Chr04 | 431 | 47.18 | 6.39 | endoplasmic reticulum |
|  | PcaHSP70-4 | Pca_137_5.64 | Chr05 | 650 | 71.33 | 5.42 | cytoplasm |
|  | PcaHSP70-5 | Pca_149_4.62 | Chr06 | 500 | 54.06 | 5.77 | plasma membrane |
|  | PcaHSP70-6 | Pca_16_40.11 | Chr07 | 550 | 60.67 | 5.66 | cytoplasm |
|  | PcaHSP70-7 | Pca_16_40.9 | Chr07 | 550 | 60.67 | 5.66 | cytoplasm |
|  | PcaHSP70-8 | Pca_100_11.48 | Chr07 | 416 | 45.42 | 5.45 | extracellular |
|  | PcaHSP70-9 | Pca_66_7.15 | Chr07 | 332 | 36.40 | 5.29 | cytoplasm |
|  | PcaHSP70-10 | Pca_66_18.49 | Chr07 | 672 | 73.33 | 4.94 | extracellular |
|  | PcaHSP70-11 | Pca_52_9.49 | Chr13 | 669 | 73.99 | 4.95 | endoplasmic reticulum |
|  | PcaHSP70-12 | Pca_96_5.36 | Chr14 | 992 | 11.26 | 5.07 | endoplasmic reticulum |
|  | PcaHSP70-13 | Pca_384_0.6 | Pca_384 | 601 | 67.24 | 7.66 | cytoplasm |
| HSP60 | PcaHSP60-1 | Pca_58_18.48 | Chr04 | 574 | 61.04 | 5.47 | mitochondrion |
| HSP40 | PcaHSP40-1 | Pca_139_4.55 | Chr03 | 174 | 19.08 | 5.31 | mitochondrion |
|  | PcaHSP40-2 | Pca_83_4.42 | Chr03 | 337 | 40.29 | 9.22 | plasma membrane |
|  | PcaHSP40-3 | Pca_2_75.25 | Chr03 | 392 | 42.74 | 8.97 | mitochondrion |
|  | PcaHSP40-4 | Pca_24_44.13 | Chr07 | 195 | 22.05 | 6.23 | cytoplasm |
|  | PcaHSP40-5 | Pca_230_0.21 | Chr07 | 358 | 40.51 | 5.33 | endoplasmic reticulum |
|  | PcaHSP40-6 | Pca_43_30.66 | Chr07 | 377 | 43.22 | 8.37 | endoplasmic reticulum |
|  | PcaHSP40-7 | Pca_21_9.15 | Chr08 | 265 | 30.71 | 9.43 | extracellular |
|  | PcaHSP40-8 | Pca_34_11.24 | Chr09 | 409 | 45.25 | 5.84 | cytoplasm |
|  | PcaHSP40-9 | Pca_46_9.37 | Chr10 | 317 | 35.87 | 8.60 | cytoplasm |
|  | PcaHSP40-10 | Pca_150_3.12 | Chr11 | 262 | 30.41 | 9.81 | mitochondrion |
|  | PcaHSP40-11 | Pca_27_38.6 | Chr12 | 353 | 38.61 | 9.04 | nucleus |
|  | PcaHSP40-12 | Pca_101_4.48 | Chr14 | 506 | 57.13 | 9.30 | nucleus |
|  | PcaHSP40-13 | Pca_1501_0.6 | Pca_1501 | 401 | 44.94 | 6.38 | cytoplasm |
| HSP20 | PcaHSP20-1 | Pca_263_0.7 | Chr03 | 225 | 25.45 | 5.75 | cytoplasm |
|  | PcaHSP20-2 | Pca_41_5.29 | Chr03 | 327 | 37.19 | 6.74 | cytoplasm |
|  | PcaHSP20-3 | Pca_2_74.33 | Chr03 | 207 | 23.72 | 7.75 | mitochondrion |
|  | PcaHSP20-4 | Pca_8_71.81 | Chr10 | 146 | 16.55 | 9.98 | mitochondrion |
|  | PcaHSP20-5 | Pca_8_37.45 | Chr10 | 164 | 18.70 | 5.60 | cytoplasm_nucleus |
|  | PcaHSP20-6 | Pca_8_37.41 | Chr10 | 135 | 16.06 | 6.21 | cytoplasm |
|  | PcaHSP20-7 | Pca_8_37.43 | Chr10 | 150 | 17.50 | 6.11 | cytoplasm_nucleus |
|  | PcaHSP20-8 | Pca_8_37.44 | Chr10 | 164 | 18.74 | 6.15 | cytoplasm_nucleus |
|  | PcaHSP20-9 | Pca_8_37.40 | Chr10 | 164 | 18.97 | 6.84 | cytoplasm_nucleus |
|  | PcaHSP20-10 | Pca_6_52.38 | Chr13 | 229 | 25.88 | 6.10 | cytoplasm_nucleus |
|  | PcaHSP20-11 | Pca_1622_0.3 | Pca_1622 | 281 | 30.17 | 6.00 | nucleus |
| HSP10 | PcaHSP10-1 | Pca_58_18.42 | Chr04 | 102 | 11.34 | 7.95 | cytoplasm |

**Fig. 1.**
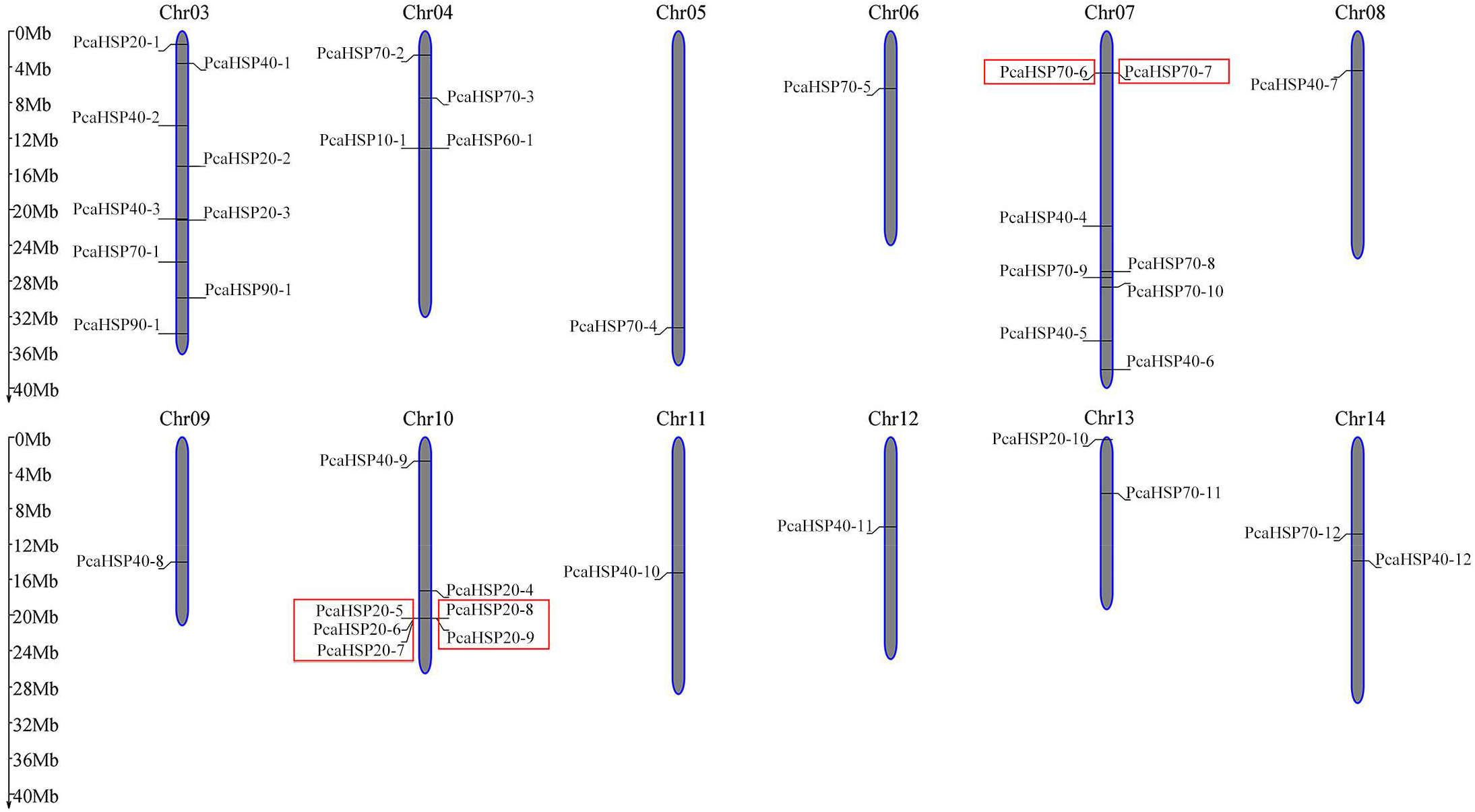
Chromosome position and tandem duplication of the HSPs in *P. canaliculata* (PcaHSP). Gray bars represent the chromosomes with the scales of chromosome length (Mb) on the left. The PcaHSPs were renamed according their position on the chromosomes. PcaHSP genes in red boxes are groups of tandem duplication.

We found two groups of tandem duplications among the PcaHSPs based on analyzing the gene duplication events (Fig. 1). One group was the paralogous gene pair PcaHSP70-6 and PcaHSP70-7 on chr07, and the other group was the HSP20 genes on chr10 (Fig. 1). We did not find any segmental duplication events in the PcaHSPs.

### 3.3 Phylogenetic analysis

Both the NJ and ML trees showed close phylogenetic relationships of each HSP family among the four species with strong support values (Fig. 2; Appendix S2: Fig. S1). The families of HSP90, HSP70, HSP60, and HSP20 were monophyletic, while the HSP40s formed a paraphyletic group as revealed from the NJ tree (Fig. 2). The phylogenetic relationships among the HSPs confirmed their classification of families, which was assigned through the Pfam database.

**Fig. 2.**
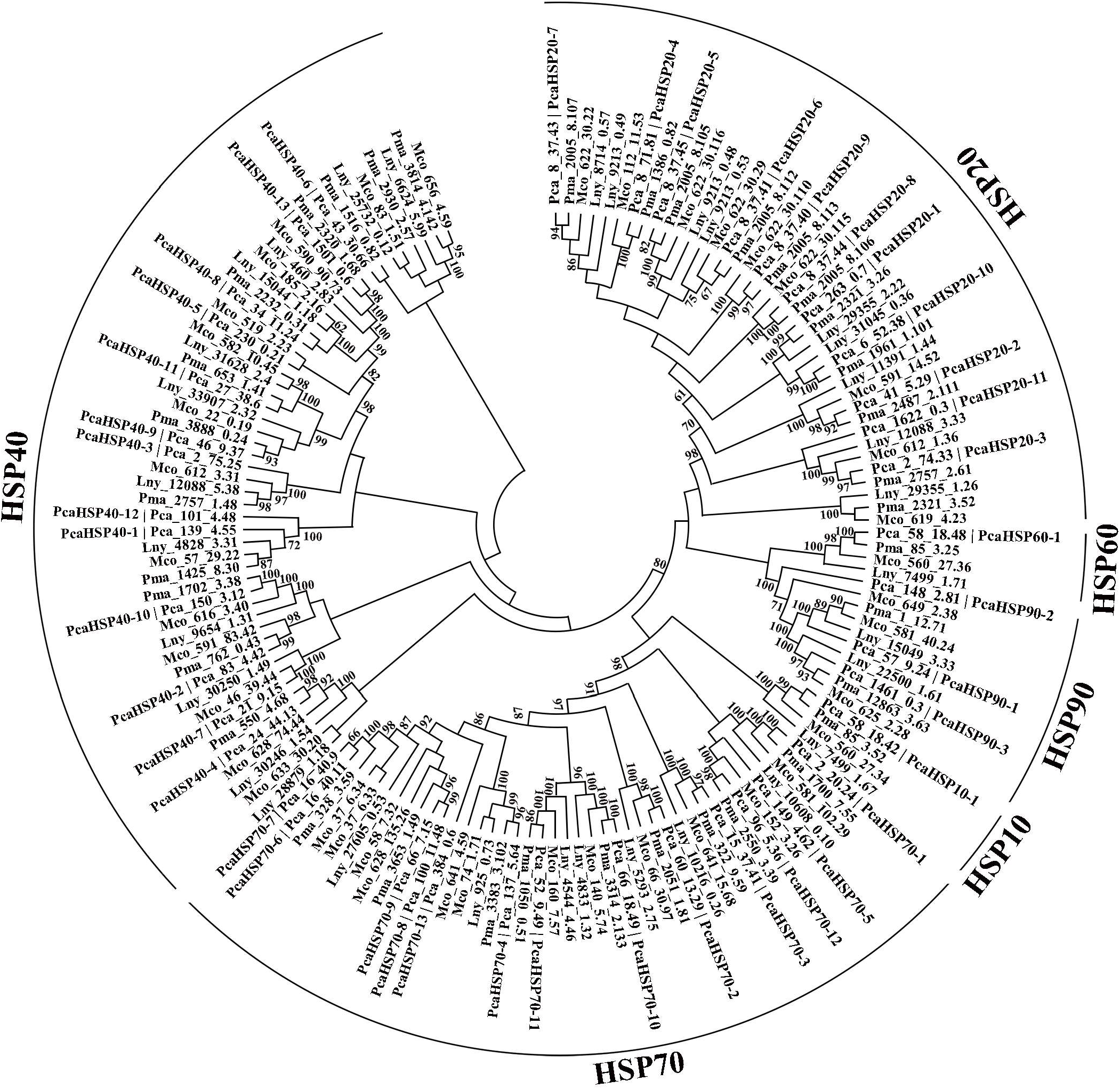
The neighbor-joining phylogenetic tree based on the HSP proteins from *P. canaliculata, P. maculata, L. nyassanus* and *M. cornuarietis*. Bootstrap values > 60% are shown.

### 3.4 Characteristics of the PcaHSPs

The sequence lengths of the PcaHSP protein varied from 102 to 992 amino acids, while the molecular weights (MWs) and isoelectric points (pIs) were in the ranges of 11.26–95.90 kDa and 4.94–9.98, respectively (Table 2). Subcellular localization analysis indicated that sixteen PcaHSPs were in the cytoplasm, seven in the mitochondrion, five in the cytoplasm_nucleus, and five in the endoplasmic reticulum (Table 2). Other PcaHSPs were located in the nucleus, extracellular space, or plasma membrane (Table 2).

Each family of PcanHSPs was classified as monophyletic according to the unrooted NJ tree based on PcanHSP protein sequences (Fig. 3). Gene structure analysis showed high divergence of the intron/exon arrangements among the families of PcaHSP (Fig. 3). Notable variation of the intron numbers was found in the PcaHSP70 genes with no introns in PcaHSP70-6, PcaHSP70-7, and PcaHSP70-9 and one intron in PcaHSP70-8, while others had multiple introns with the maximum found in PcaHSP70-12. For the introns in the PcaHSP20s, there were none in PcaHSP20-2 and one in PcaHSP20-11, and the others were more similar to two introns, except PcaHSP20-1, which had four introns. All the PcaHSP90 family genes have eight introns except PcaHSP90-2, which contains nine introns. For the PcaHSP40s, no introns were found in PcaHSP40-10 and PcaHSP40-11, while other PcaHSP40s had introns varying from four to eleven.

**Fig. 3.**
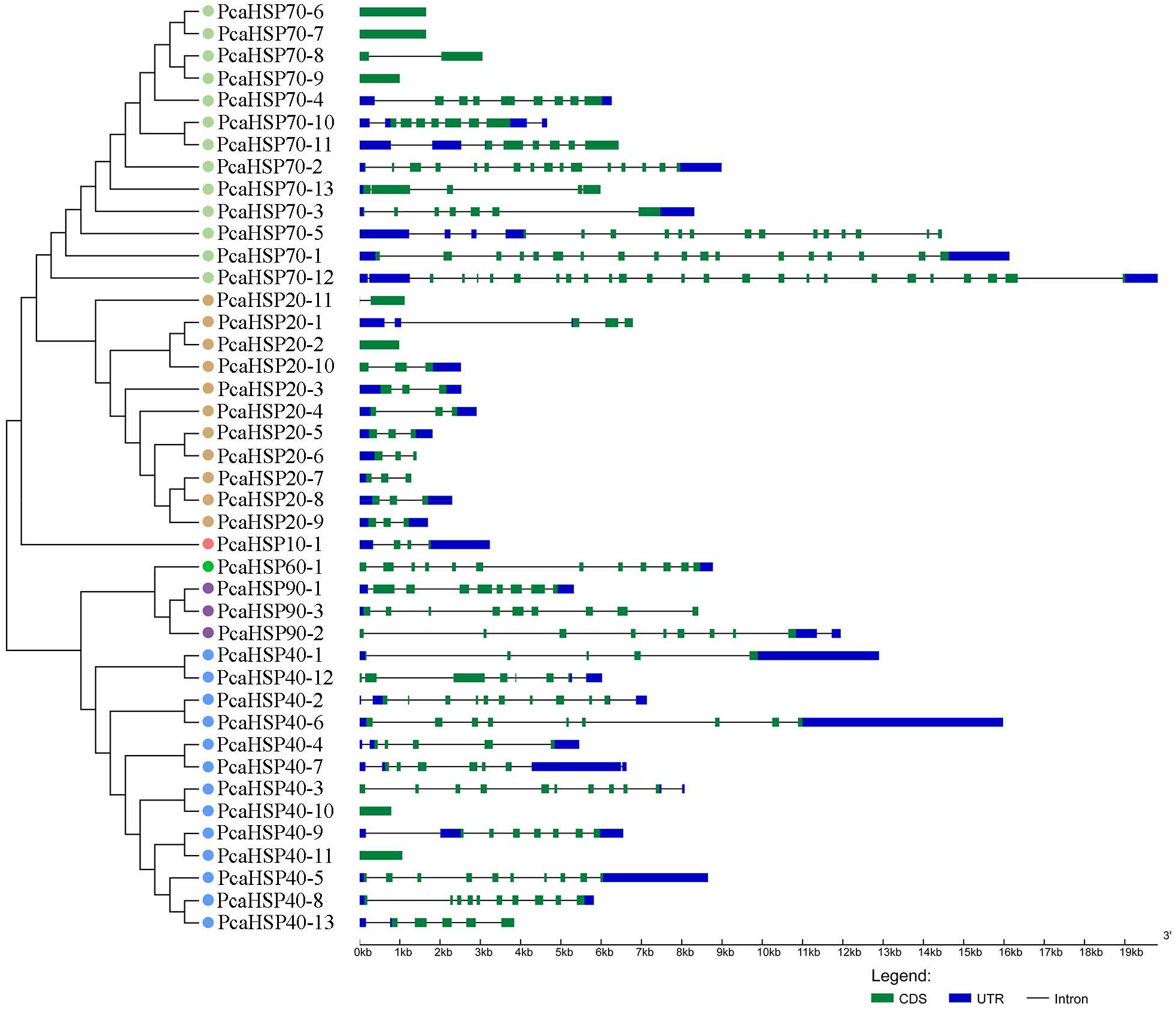
Phylogenetic relationships and gene structures of the HSPs in *P. canaliculate* (PcaHSP). CDS, coding sequence for protein; UTR, untranslated region

A total of 20 putative motifs were predicted from the 42 PcaHSP proteins (Fig. 4; Appendix S2: Fig. S2). Most motifs were found in PcaHSP70, while the fewest motifs were found in PcaHSP10 (Fig. 4). The composition and arrangement of motifs in each HSP family were conserved, whereas considerable diversity was observed among families (Fig. 4). Motifs 1, 4, 6, 8, and 12 were shared by at least two HSP families, and the other motifs were family-specific (Fig. 4).

**Fig. 4.**
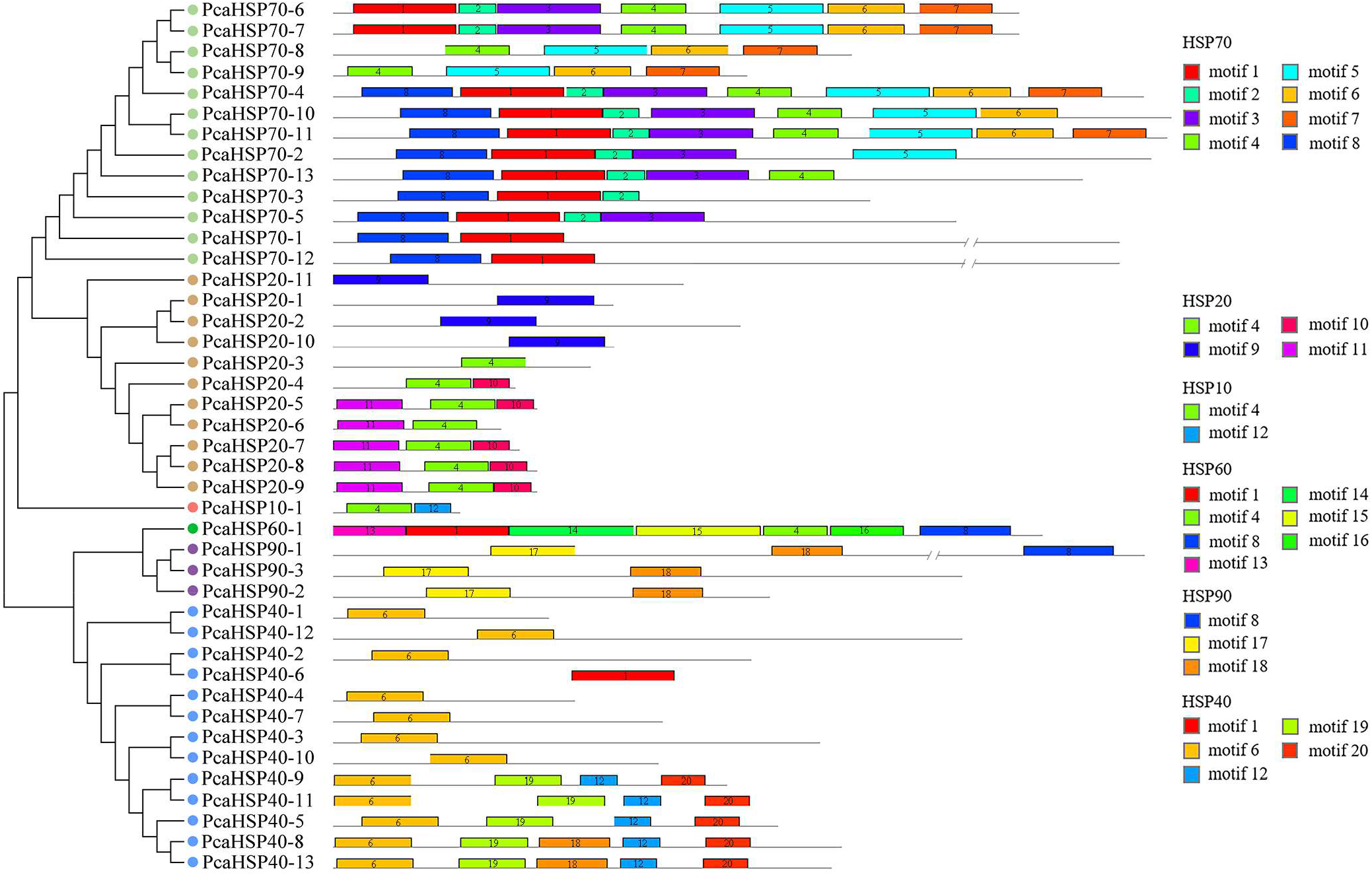
Phylogenetic relationships and conserved motifs of the HSPs in *P. canaliculata* (PcaHSP). Each motif is represented by a colored box with numbers. The numbers shown on each gene correspond to the motif numbers. Lengths of motifs for each PcaHSP protein are shown proportionally.

### 3.5 Expression profiles of the PcaHSPs in different tissues

We employed RNA-seq data to examine the expression patterns of PcaHSPs in different tissues of *P. canaliculata* (Fig. 5a). The PcaHSPs revealed similar expression patterns in the hepatopancreas, kidney, gill, ovary, and testis, with several exceptions. For instance, PcaHSP90-1, PcaHSP90-3, PcaHSP70-4, PcaHSP70-11, and PcaHSP60-1 were among the genes with high expression levels, but PcanHSP70-10, PcanHSP40-10, PcanHSP40-12, and PcanHSP20-10 were expressed at low levels in all five tissues (Fig. 5a). The members of the PcaHSP90, PcaHSP60 and PcaHSP10 families were among the highly expressed HPSs in different tissues, while genes in the families of PcaHSP70, PcaHSP40, and PcaHSP20 varied in expression levels (Fig. 5a). In particular, several genes of PcaHSP20 revealed tissue-specific expression levels; for example, PcaHSP20-4, PcaHSP20-5, and PcaHSP20-8 were expressed at extremely high levels in the hepatopancreas but at low levels in other tissues; PcaHSP20-9 displayed a higher expression level in the ovary, and PcaHSP20-1 and PcaHSP20-2 were expressed at higher levels in the testis than in other tissues (Fig. 5a).

**Fig. 5.**
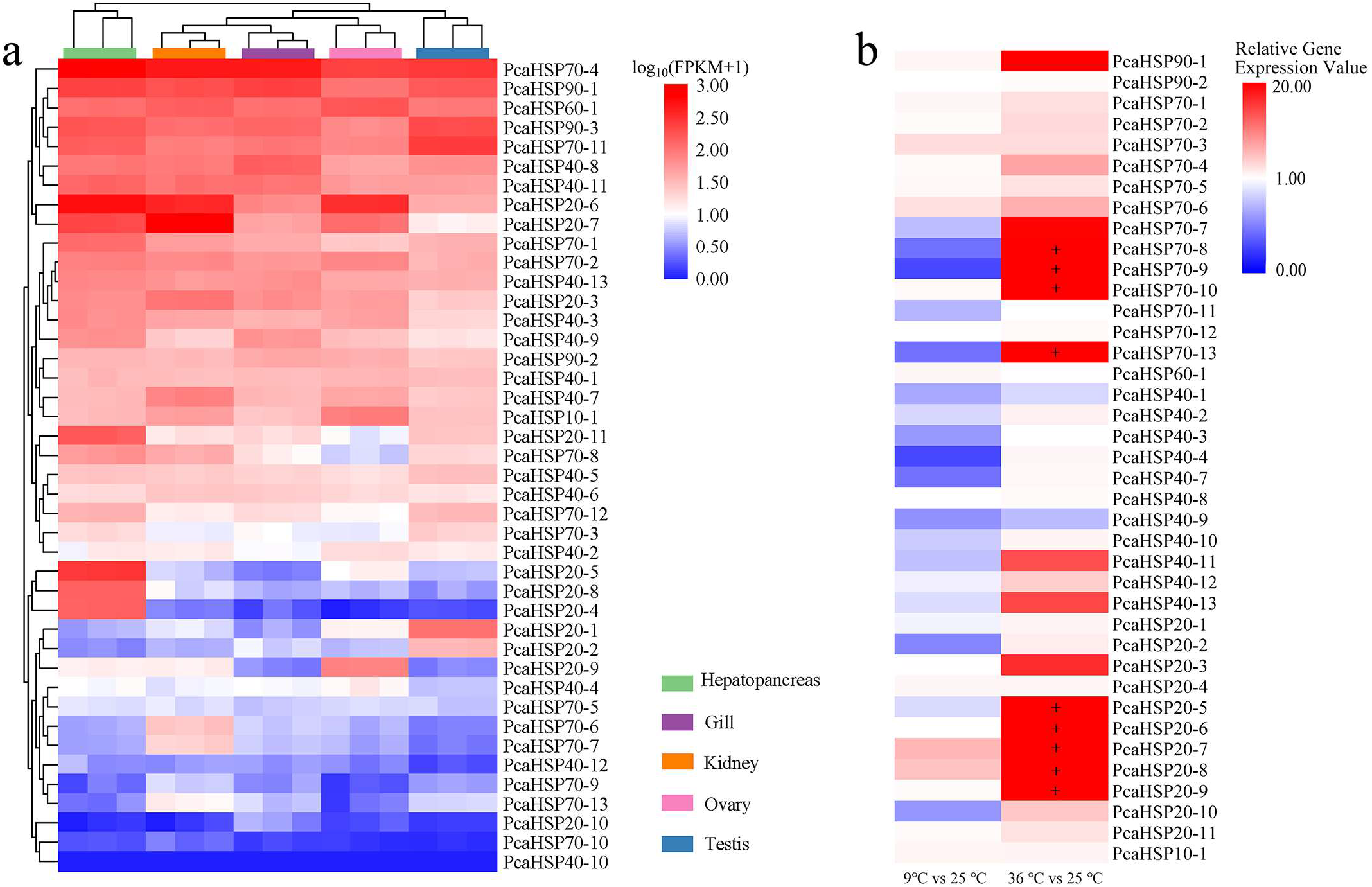
Heatmap of the expression profiles of the HSPs in *P. canaliculata* (PcaHSP) in different tissues according to the RNA-seq data (a) and their expression patterns under hot and cold stresses (b). 9°vs25° represents the cold stress at 9° for 2 h following 25° for 1 h vs. 25° for 3 h; 36°vs25° represents the hot stress at 36° for 2 h following 25°for 1 h vs. 25° for 3 h. Relative gene expression values were calculated using the 2^−^ △△Ct method; “+” meant that the value was greater than 20.

### 3.6 Gene expression of PcaHSPs under temperature stress

A majority of the 42 PcaHSPs can be amplified to produce specific target bands with our designed primers (Appendix S2: Table S3). However, we were not able to amplify PcaHSP90-3, PcaHSP40-5, and PcaHSP40-6, although we tried three pairs of primers for each gene.

Under high-temperature treatment, 36 out of the 39 HSP genes were upregulated, and three HSP40s (PcaHSP40-1, PcaHSP40-3, and PcaHSP40-9) were downregulated (Fig. 5b). Most of the PcaHSP20 and PcaHSP70 genes were heat-induced (Fig. 5b). Notably, the relative expression levels of four HSP70s (PcaHSP70-8–PcaHSP70-10, PcaHSP70-13) and five HSP20s (PcaHSP20-5–PcaHSP20-9) were extremely highly upregulated, with fold changes in the range of 27.6-833.9, compared with the normal condition under high-temperature treatment.

Under low-temperature treatment, 20 HSP genes were induced, and 19 HSP genes were downregulated. Thirteen HSPs, including PcaHSP90-1, PcaHSP70-3, PcaHSP70-6, PcaHSP20-7, and PcaHSP20-8, displayed mild upregulation under low-temperature treatment; despite being lower than 20-fold, PcaHSP20-7 and PcaHSP20-8 were the two genes with the highest expression levels compared with the others (Fig. 5b).

Nineteen genes, including PcaHSP70-8, PcaHSP70-9, PcaHSP70-13 PcaHSP40-4, PcaHSP40-7, PcaHSP20-2, and PcaHSP20-10, were downregulated under low temperature (Fig. 5b).

## 4. Discussion

The HSP proteins play key roles in assisting the new protein folding as chaperones to respond to environmental stress, prevent unwanted conformation or aggregation, and transport proteins across membranes in cells [41, 42]. Previous studies have shown that HSP proteins participate in various physiological activities in *P. canaliculata*, including immunity [43], stress responses [38, 44, 45], and the activity-estivation cycle [46]. The recent release of whole genomic data of four apple snail species, i.e., *P. canaliculata, P. maculata, L. nyassanus*, and *M. cornuarietis*, has enabled an integrated identification of the crucial stress adaptation-related gene families using a comparative genomics strategy [15]. Previously, the GST family, which plays a vital role in the regulation of detoxification and redox balance of ROS, was characterized in *P. canaliculata*, which revealed 30 GST genes belonging to seven classes [47]. Our study represents the most comprehensive study of the HSP superfamily in apple snails.

We found 32 to 42 HSP genes encoding six HSP families from the genomes of *P. canaliculata, P. maculata, L. nyassanus*, and *M. cornuarietis*. Although the genome sizes of *M. cornuarietis* and *L. nyassanus* are larger than those of the *Pomacea* species, the maximum numbers of HSP genes were characterized from *P. canaliculata*. HSP70, HSP40, and HSP20 were abundant in apple snails, indicating that the three families were expanded. Gene duplication events, such as segmental duplication, whole genome duplication, and tandem duplication, often play crucial roles in the expansion and evolution of gene families in organisms and the production of new functions [48, 49]. Analyzing the gene duplication events of the PcaHSPs based on multiple collinear scanning has demonstrated that tandem duplications occurred in both the HSP70 and HSP20 families. The HSP70 genes were found to undergo extensive expansion and were mainly attributed to tandem duplication events in the yesso scallop *Patinopecten yessoensis* [50]. Our results confirmed that tandem duplication played an indispensable role in the expansion of the HSP family in apple snails.

The cloning and expression of HSPs has only been studied in *P. canaliculata* among apple snails due to its great economic importance as a highly invasive species. Previously, more than twenty HSP genes were identified from *P. canaliculata* after different stimuli, which involved complete or nearly complete members of the HSP90, HSP70, and HSP40 families [38]. However, most of the previous studies focused on one or several HSP genes in *P. canaliculata*: HSP70 and HSP90 have been the most extensively studied [10, 26, 27], followed by HSP40 [28] and HSP60 [10]. Comparing the protein sequences of the published HSP genes with the PcaHSP families identified in this study, we confirmed the HSC70 reported by Zheng [27] to be PcaHSP70-4, the HSP90, HSP70, and HSP60 characterized by Xu [10] to be PcaHSP90-1, PcaHSP70-8, and PcaHSP60-1, respectively, and the HSP40 characterized by Xu [28] to be PcaHSP40-9 (Appendix S2: Fig. S3). The responses to temperature stress of the abovementioned genes in previous studies have indicated similar patterns to the results in our study.

We found that the HSP genes in the same family shared a close phylogenetic relationship and a conserved motif composition in *P. canaliculata*. Each HSP family has a specific motif that may be crucial for their functions. However, the intron patterns in each HSP family of apple snails varied greatly. The introns in eukaryotic genomes that split gene structures are considered crucial to the origin and diversification of organisms [51]. The number, position, size, and sequence context are the four parameters used to characterize introns [52]. Compared with the number and position of introns, the sequence and size of introns are usually more variable [53]. In the HSP families of *P. canaliculata*, we observed considerable variation in not only the intron size and sequence. Moreover, we observed several intronless HSP genes, including PcaHSP70-6, PcaHSP70-7, and PcaHSP70-9. Muliple intronless genes of HSP70s were also found from the genome-wide characterization of the HSP family of the invasive *Bemisia tabaci* [31]. Intron gain and loss events over a dynamic evolutionary history are the most predominant hypothesis for explaining intron variation in eukaryotes, although debates still exist about either gains or losses that are preferential in different taxa [54, 55].

sHSPs play multiple important roles in physiological and pathophysiological processes and stress tolerance with either broad or restricted tissue distribution; for instance, HSP20 is ubiquitously distributed but abundant in muscle in humans [56]. Our analyses of the expression profiles of *P. canaliculata* indicated that most HSP families have similar expression patterns in different tissues. However, a couple of HSPs, mainly from the PcaHSP20 family, showed tissue-specific expression levels, which indicated a possible tissue-specific distribution. Knowledge about the molecular mechanism involved in temperature stress tolerance is important not only to basic science but also to applied biotechnology. A large number of studies have shown that HSP proteins are related to heat and cold stress. In particular, HSP70, which often serves as a sensitive indicator of the heat shock response, was applied to reflect the thermal senilities of species [57]. Our results showed that most PcaHSP70 was induced after heat shock despite various changes in expression levels. Meanwhile, nearly all the PcaHSP20 were heat shock induced, with several of them being extremely upregulated in the hepatopancreas. To our knowledge, the sHSP family has not been investigated in *P. canaliculata*. Considering the tissue specificity, the temperature tolerance patterns of PcaHSP20 in other tissues still need further investigation. Under genome-wide analysis, many species showed more HSP genes in response to cold treatment than to thermal treatment [31, 58]. However, compared with heat stress, there were fewer PcaHSPs involved in the response to cold stress and with lower expression levels.

In summary, this study identified a full set of HSP family members from the genomes of four apple snails and characterized the HSPs of the invasive species *P. canaliculata*. We renamed the PcaHSPs based on their positions on chromosomes and conducted integrated analyses of their physical and chemical properties, including gene structure, phylogenetic relationships, chromosome distribution, tissue-expression specificity, and temperature stress responses. Our results have provided basic information for further research on the molecular function of HSPs in adaptability to diverse environments and will help to understand the molecular mechanism of the invasive success of apple snails.

## Acknowledgements

This work was supported by grants from the National Natural Science Foundation of China (32171668), the Fundamental Research Funds for the Provincial Universities of Zhejiang (2021YW06), Promoting Scientific Research Cooperation with Canada, Australia, New Zealand, and Latin America and High-level Talent Training Projects of the Chinese Scholarship Council to Q.-Q. Yang.

## Supporting information

Appendix S1 Sequence list of the HSP genes identified from the genomes of *P. canaliculata, P. maculata, L. nyassanus*, and *M. cornuarietis*

Appendix S2 Supplementary tables and figures.

## References

[1] M.G. Santoro, Heat shock factors and the control of the stress response, Biochem. Pharmacol. 59 (2000) 55–63. https://doi.org/10.1016/s0006-2952(99)00299-3.

[2] P. Srivastava, Roles of heat-shock proteins in innate and adaptive immunity, Nat. Rev. Immunol. 2 (2002) 185–194. https://doi.org/10.1038/nri749.

[3] R. Arya, M. Mallik, S.C. Lakhotia, Heat shock genes - integrating cell survival and death, J. Biosci. 32 (2007) 595–610. https://doi.org/10.1007/s12038-007-0059-3.

[4] M.E. Feder, G.E. Hofmann, Heat-shock proteins, molecular chaperones, and the stress response: evolutionary and ecological physiology, Annu. Rev. Physiol. 61 (1999) 243–282. https://doi.org/10.1146/annurev.physiol.61.1.243.

[5] N. Kourtis, N. Tavernarakis, Small heat shock proteins and neurodegeneration: recent developments, Biomol. Concepts 9 (2018) 94–102. https://doi.org/10.1515/bmc-2018-0009.

[6] M. Zolkiewski, T. Zhang, M. Nagy, Aggregate reactivation mediated by the Hsp100 chaperones, Arch. Biochem. Biophys. 520 (2012) 1–6. https://doi.org/10.1016/j.abb.2012.01.012.

[7] M. Haslbeck, E. Vierling, A first line of stress defense: small heat shock proteins and their function in protein homeostasis, J. Mol. Biol. 427 (2015) 1537–1548. https://doi.org/10.1016/j.jmb.2015.02.002.

[8] M. Haslbeck, S. Weinkauf, J. Buchner, Small heat shock proteins: simplicity meets complexity, J. Biol. Chem. 294 (2019) 2121–2132. https://doi.org/10.1074/jbc.REV118.002809.

[9] J.G. Kiang, G.C. Tsokos, Heat shock protein 70 kDa: molecular biology, biochemistry, and physiology, Pharmacol. Ther. 80 (1998) 183–201. https://doi.org/10.1016/s0163-7258(98)00028-x.

[10] Y. Xu, G. Zheng, S. Dong, G. Liu, X. Yu, Molecular cloning, characterization and expression analysis of HSP60, HSP70 and HSP90 in the golden apple snail, Pomacea canaliculata, Fish Shellfish Immunol. 41 (2014) 643–653. https://doi.org/10.1016/j.fsi.2014.10.013.

[11] M.K. Nelson, B.C. Cruz, K.L. Buena, H. Nguyen, J.T. Sullivan, Effects of abnormal temperature and starvation on the internal defense system of the schistosome-transmitting snail Biomphalaria glabrata, J. Invertebr. Pathol. 138 (2016) 18–23. https://doi.org/10.1016/j.jip.2016.05.009.

[12] Y. Huang, X. Cai, Z. Zou, S. Wang, G. Wang, Y. Wang, Z. Zhang, Molecular cloning, characterization and expression analysis of three heat shock responsive genes from Haliotis diversicolor, Fish Shellfish Immunol. 36 (2014) 590–599. https://doi.org/10.1016/j.fsi.2013.11.013.

[13] K.A. Hayes, R.L. Burks, A. Castro-Vazquez, P.C. Darby, H. Heras, P.R. Martin, J.W. Qiu, S.C. Thiengo, I.A. Vega, T. Wada, Y. Yusa, S. Burela, M.P. Cadierno, J.A. Cueto, F.A. Dellagnola, M.S. Dreon, M.V. Frassa, M. Giraud-Billoud, M.S. Godoy, S. Ituarte, E. Koch, K. Matsukura, M.Y. Pasquevich, C. Rodriguez, L. Saveanu, M.E. Seuffert, E.E. Strong, J. Sun, N.E. Tamburi, M.J. Tiecher, R.L. Turner, P.L. Valentine-Darby, R.H. Cowie, Insights from an integrated view of the biology of apple snails (Caenogastropoda: Ampullariidae), Malacologia 58 (2015) 245–302. https://doi.org/10.4002/040.058.0209.

[14] K. Hayes, R. Cowie, A. Jørgensen, R. Schultheiß, C. Albrecht, S. Thiengo, Molluscan models in evolutionary biology: apple snails (Gastropoda: Ampullariidae) as a system for addressing fundamental questions, Am. Malacol. Bull. 27 (2009) 47–58. https://doi.org/10.4003/006.027.0204.

[15] J. Sun, H. Mu, J.C.H. Ip, R. Li, T. Xu, A. Accorsi, A. Sánchez Alvarado, E. Ross, Y. Lan, Y. Sun Castro-Vazquez, I.A. Vega, H. Heras, S. Ituarte, B. Van Bocxlaer, K.A. Hayes, R.H. Cowie, Z. Zhao, Y. Zhang, P.Y. Qian, J.W. Qiu, Signatures of divergence, invasiveness, and terrestrialization revealed by four apple snail genomes, Mol. Biol. Evol. 36 (2019) 1507–1520. https://doi.org/10.1093/molbev/msz084.

[16] K.A. Hayes, R.H. Cowie, S.C. Thiengo, E.E. Strong, Comparing apples with apples: clarifying the identities of two highly invasive neotropical Ampullariidae (Caenogastropoda), Zool. J. Linn. Soc. 166 (2012) 723–753. https://doi.org/10.1111/j.1096-3642.2012.00867.x.

[17] K. Matsukura, K.A. Hayes, R.H. Cowie, Eleven microsatellite loci for two invasive freshwater apple snails, Pomacea canaliculata and P. maculata (Ampullariidae), Biol. Invasions 18 (2016) 3397–3400. https://doi.org/10.1007/s10530-016-1237-8.

[18] M.E. Seuffert, P.R. Martín, Juvenile growth and survival of the apple snail Pomacea canaliculata (Caenogastropoda: Ampullariidae) reared at different constant temperatures, Springerplus 2 (2013) 312. https://doi.org/10.1186/2193-1801-2-312.

[19] M.E. Seuffert, L. Saveanu, P.R. Martín, Threshold temperatures and degree-day estimates for embryonic development of the invasive apple snail Pomacea canaliculata (Caenogastropoda: Ampullariidae), Malacologia 55 (2012) 209–217. https://doi.org/10.4002/040.055.0203.

[20] A. Estebenet, P. Martín, Pomacea canaliculata (Gastropoda: Ampullariidae): life-history traits and their plasticity, Biocell 26 (2002) 83–89. PMID: 12058384.

[21] K. Matsukura, M. Okuda, N.J. Cazzaniga, T. Wada, Genetic exchange between two freshwater apple snails, Pomacea canaliculata and Pomacea maculata invading East and Southeast Asia, Biol. Invasions 15 (2013) 2039–2048. https://doi.org/10.1007/s10530-013-0431-1.

[22] K. Matsukura, H. Tsumuki, Y. Izumi, T. Wada, Changes in chemical components in the freshwater apple snail, Pomacea canaliculata (Gastropoda: Ampullariidae), in relation to the development of its cold hardiness, Cryobiology 56 (2008) 131–137. https://doi.org/10.1016/j.cryobiol.2007.12.001.

[23] K. Heiler, P. Viktor, V. Oheimb, K. Lemens, C. Albrecht, Studies on the temperature dependence of activity and on the diurnal activity rhythm of the invasive Pomacea canaliculata (Gastropoda: Ampullariidae), Mollusca 26 (2008) 73–81.

[24] T. Wada, K. Yoshida, Burrowing by the apple snail, Pomacea canaliculata (Lamarck); daily periodicity and factors affecting burrowing, Kyushu Byogaichu Kenkyukaiho 46 (2000) 88–93. https://doi.org/10.4241/kyubyochu.46.88.

[25] A. Stevens, Z. Welch, P. Darby, H. Percival, Temperature effects on florida apple snail activity: implications for snail kite foraging success and distribution, Wildl. Soc. Bull. 30 (2002) 75–81. https://doi.org/10.2307/3784638.

[26] H.M. Song, X.D. Mu, D.E. Gu, D. Luo, Y.X. Yang, M. Xu, J.R. Luo, J.E. Zhang, Y.C. Hu, Molecular characteristics of the HSP70 gene and its differential expression in female and male golden apple snails (Pomacea canaliculata) under temperature stimulation, Cell Stress Chaperones 19 (2014) 579–589. https://doi.org/10.1007/s12192-013-0485-0.

[27] G. Zheng, S. Dong, Y. Hou, K. Yang, X. Yu, Molecular characteristics of HSC70 gene and its expression in the golden apple snails, Pomacea canaliculata (Mollusca: Gastropoda), Aquaculture 358 (2012) 41–49. https://doi.org/10.1016/j.aquaculture.2012.06.002.

[28] Y. Xu, G. Zheng, G. Liu, Q. Yang, X. Yu, Molecular cloning, characterization of Pomacea canaliculata HSP40 and its expression analysis under temperature change, J. Therm. Biol. 81 (2019) 59–65. https://doi.org/10.1016/j.jtherbio.2019.02.006.

[29] X. Huang, S. Li, Y. Gao, A. Zhan, Genome-wide identification, characterization and expression analyses of heat shock protein-related genes in a highly invasive ascidian Ciona savignyi, Front. Physiol. 9 (2018) 1043. https://doi.org/10.3389/fphys.2018.01043.

[30] F.L. Si, L. Qiao, Q.Y. He, Y. Zhou, Z.T. Yan, B. Chen, HSP superfamily of genes in the malaria vector Anopheles sinensis: diversity, phylogenetics and association with pyrethroid resistance, Malar. J. 18 (2019) 132. https://doi.org/10.1186/s12936-019-2770-6.

[31] X.R. Wang, C. Wang, F.X. Ban, D.T. Zhu, S.S. Liu, X.W. Wang, Genome-wide identification and characterization of HSP gene superfamily in whitefly (Bemisia tabaci) and expression profiling analysis under temperature stress, Insect Sci. 26 (2019) 44–57. https://doi.org/10.1111/1744-7917.12505.

[32] P.K. Meher, T.K. Sahu, S. Gahoi, A.R. Rao, ir-HSP: improved recognition of heat shock proteins, their families and sub-types based on g-spaced dipeptide features and support vector machine, Front. Genet. 8 (2017) 235. https://doi.org/10.3389/fgene.2017.00235.

[33] J. Chao, K. Yingzhen, Q. Wang, S. Yuhe, G. Daping, J. Lv, L. Guanshan, MapGene2Chrom, a tool to draw gene physical map based on Perl and SVG languages, Hereditas (Beijing) 37 (2015) 91–97. https://doi.org/10.16288/j.yczz.2015.01.013.

[34] Y. Wang, H. Tang, J.D. Debarry, X. Tan, J. Li, X. Wang, T.H. Lee, H. Jin, B. Marler, H. Guo, J.C. Kissinger, A.H. Paterson, MCScanX: a toolkit for detection and evolutionary analysis of gene synteny and collinearity, Nucleic Acids Res. 40 (2012) e49. https://doi.org/10.1093/nar/gkr1293.

[35] S. Kumar, G. Stecher, M. Li, C. Knyaz, K. Tamura, MEGA X: molecular evolutionary genetics analysis across computing platforms, Mol. Biol. Evol. 35 (2018) 1547–1549. https://doi.org/10.1093/molbev/msy096.

[36] L.T. Nguyen, H.A. Schmidt, A. von Haeseler, B.Q. Minh, IQ-TREE: a fast and effective stochastic algorithm for estimating maximum-likelihood phylogenies, Mol. Biol. Evol. 32 (2015) 268–274. https://doi.org/10.1093/molbev/msu300.

[37] T.L. Bailey, M. Boden, F.A. Buske, M. Frith, C.E. Grant, L. Clementi, J. Ren, W.W. Li, W.S. Noble, MEME SUITE: tools for motif discovery and searching, Nucleic Acids Res. 37 (2009) W202–W208. https://doi.org/10.1093/nar/gkp335.

[38] C. Liu, Y. Zhang, Y. Ren, H. Wang, S. Li, F. Jiang, L. Yin, X. Qiao, G. Zhang, W. Qian, B. Liu, W. Fan, The genome of the golden apple snail Pomacea canaliculata provides insight into stress tolerance and invasive adaptation, Gigascience 7 (2018) 1–13. https://doi.org/10.1093/gigascience/giy101.

[39] G.P. Wagner, K. Kin, V.J. Lynch, Measurement of mRNA abundance using RNA-seq data: RPKM measure is inconsistent among samples, Theory Biosci 131 (2012) 281–285. https://doi.org/10.1007/s12064-012-0162-3.

[40] A.I. Saeed, V. Sharov, J. White, J. Li, W. Liang, N. Bhagabati, J. Braisted, M. Klapa, T. Currier, M. Thiagarajan, A. Sturn, M. Snuffin, A. Rezantsev, D. Popov, A. Ryltsov, E. Kostukovich, I. Borisovsky, Z. Liu, A. Vinsavich, V. Trush, J. Quackenbush, TM4: a free, open-source system for microarray data management and analysis, BioTechniques 34 (2003) 374–378. https://doi.org/10.2144/03342mt01.

[41] K.C. Kregel, Heat shock proteins: modifying factors in physiological stress responses and acquired thermotolerance, J. Appl. Physiol. 92 (2002) 2177–2186. https://doi.org/10.1152/japplphysiol.01267.2001.

[42] C.J. Park, Y.S. Seo, Heat shock proteins: a review of the molecular chaperones for plant immunity, Plant Pathol. J. 31 (2015) 323–333. https://doi.org/10.5423/ppj.Rw.08.2015.0150.

[43] F. Boraldi, F.D. Lofaro, A. Accorsi, E. Ross, D. Malagoli, Toward the molecular deciphering of Pomacea canaliculata immunity: first proteomic analysis of circulating hemocytes, Proteomics 19 (2019) e1800314. https://doi.org/10.1002/pmic.201800314.

[44] F. Boraldi, F.D. Lofaro, G. Bergamini, A. Ferrari, D. Malagoli, Pomacea canaliculata ampullar proteome: a nematode-based bio-pesticide induces changes in metabolic and stress-related pathways, Biology (Basel) 10 (2021) 1049. https://doi.org/10.3390/biology10101049.

[45] S.R. Park, Y.K. Choi, H.J. Lee, S.Y. Lee, Y.K. Kim, Characterization of heat shock protein 70 in freshwater snail, Semisulcospira coreana in response to temperature and salinity, Journal of Marine Life Science 5 (2020) 17–24.

[46] M. Giraud-Billoud, I.A. Vega, M.E. Tosi, M.A. Abud,M.L. Calderón, A. Castro-Vazquez, Antioxidant and molecular chaperone defences during estivation and arousal in the South American apple snail Pomacea canaliculata, J. Exp. Biol. 216 (2013) 614–622. https://doi.org/10.1242/jeb.075655.

[47] Y. Lin, Q. Xiao, Q. Hao, Z. Qian, X. Li, P. Li, H. Li, L. Chen, Genome-wide identification and functional analysis of the glutathione S-transferase (GST) family in Pomacea canaliculata, Int. J. Biol. Macromol. 193 (2021) 2062–2069. https://doi.org/10.1016/j.ijbiomac.2021.11.038.

[48] M.R. Mehan, N.B. Freimer, R.A. Ophoff, A genome-wide survey of segmental duplications that mediate common human genetic variation of chromosomal architecture, Hum. Genomics 1 (2004) 335–344. https://doi.org/10.1186/1479-7364-1-5-335.

[49] M. Liu, W. Sun, Z. Ma, L. Huang, Q. Wu, Z. Tang, T. Bu, C. Li, H. Chen, Genome-wide identification of the SPL gene family in tartary buckwheat (Fagopyrum tataricum) and expression analysis during fruit development stages, BMC Plant Biol. 19 (2019) 299. https://doi.org/10.1186/s12870-019-1916-6.

[50] J. Cheng, X. Xun, Y. Kong, S. Wang, Z. Yang, Y. Li, D. Kong, S. Wang, L. Zhang, X. Hu, Z. Bao, Hsp70 gene expansions in the scallop Patinopecten yessoensis and their expression regulation after exposure to the toxic dinoflagellate Alexandrium catenella, Fish Shellfish Immunol. 58 (2016) 266–273. https://doi.org/10.1016/j.fsi.2016.09.009.

[51] I.B. Rogozin, L. Carmel, M. Csuros, E.V. Koonin, Origin and evolution of spliceosomal introns, Biol. Direct. 7 (2012) 11. https://doi.org/10.1186/1745-6150-7-11.

[52] B. Chen, J. Shao, H. Zhuang, J. Wen, Evolutionary dynamics of triosephosphate isomerase gene intron location pattern in Metazoa: a new perspective on intron evolution in animals, Gene 602 (2017) 24–32. https://doi.org/10.1016/j.gene.2016.11.027.

[53] I.B. Rogozin, Y.I. Wolf, A.V. Sorokin, B.G. Mirkin, E.V. Koonin, Remarkable interkingdom conservation of intron positions and massive, lineage-specific intron loss and gain in eukaryotic evolution, Curr. Biol. 13 (2003) 1512–1517. https://doi.org/10.1016/s0960-9822(03)00558-x.

[54] W.G. Qiu, N. Schisler, A. Stoltzfus, The evolutionary gain of spliceosomal introns: sequence and phase preferences, Mol. Biol. Evol. 21 (2004) 1252–1263. https://doi.org/10.1093/molbev/msh120.

[55] M.Y. Ma, J. Xia, K.X. Shu, D.K. Niu, Intron losses and gains in the nematodes, Biology Direct. 17 (2022) 13. https://doi.org/10.1186/s13062-022-00328-8.

[56] R. Bakthisaran, R. Tangirala, M. Rao Ch, Small heat shock proteins: role in cellular functions and pathology, Biochim. Biophys. Acta. 1854 (2015) 291–319. https://doi.org/10.1016/j.bbapap.2014.12.019.

[57] L. Tomanek, E. Sanford, Heat-shock protein 70 (Hsp70) as a biochemical stress indicator: an experimental field test in two congeneric intertidal gastropods (genus: Tegula), Biol. Bull. 205 (2003) 276–284. https://doi.org/10.2307/1543291.

[58] F. Jiang, G. Chang, Z. Li, M. Abouzaid, X. Du, J.J. Hull, W. Ma, Y. Lin, The HSP/co-chaperone network in environmental cold adaptation of Chilo suppressalis, Int. J. Biol. Macromol. 187 (2021) 780–788. https://doi.org/10.1016/j.ijbiomac.2021.07.113.

