## Appendix S2 for "Genome-wide identification and characterization of the HSP gene superfamily in apple snails (Gastropoda: Ampullariidae) and expression analysis under temperature stress"

Table S1 List of species and GenBank accession numbers used in phylogenetic analyses of HSPs in apple snails

| Sequence_ID | Predicted_class_of_HSP | Species |
| --- | --- | --- |
| XP_012940482.1 | HSP90 | *Aplysia californica* |
| XP_005102885.1 | HSP90 | *Aplysia californica* |
| XP_012945462.1 | HSP90 | *Aplysia californica* |
| XP_013072750.1 | HSP90 | *Biomphalaria glabrata* |
| ACX94847.1 | HSP90 | *Haliotis discus hannai* |
| XP_009065664.1 | HSP90 | *Lottia gigantea* |
| XP_009055707.1 | HSP90 | *Lottia gigantea* |
| XP_009045629.1 | HSP90 | *Lottia gigantea* |
| AIZ03410.1 | HSP90 | *Pomacea canaliculata* |
| AYH91699.1 | HSP90 | *Pomacea canaliculata* |
| XP_025107756.1 | HSP90 | *Pomacea canaliculata* |
| XP_025084764.1 | HSP90 | *Pomacea canaliculata* |
| PVD35258.1 | HSP90 | *Pomacea canaliculata* |
| PVD24179.1 | HSP90 | *Pomacea canaliculata* |
| PVD34873.1 | HSP90 | *Pomacea canaliculata* |
| XP_005097387.1 | HSP60 | *Aplysia californica* |
| ACL00842.1 | HSP60 | *Biomphalaria glabrata* |
| XP_009045001.1 | HSP60 | *Lottia gigantea* |
| AIZ03411.1 | HSP60 | *Pomacea canaliculata* |
| CAA78757.1 | HSP70 | *Aplysia californica* |
| CAA78755.1 | HSP70 | *Aplysia californica* |
| XP_005103834.1 | HSP70 | *Aplysia californica* |
| XP_005100356.1 | HSP70 | *Aplysia californica* |
| XP_005100354.1 | HSP70 | *Aplysia californica* |
| XP_005100353.1 | HSP70 | *Aplysia californica* |
| XP_005102694.1 | HSP70 | *Aplysia californica* |
| XP_005098078.1 | HSP70 | *Aplysia californica* |
| XP_005098119.1 | HSP70 | *Aplysia californica* |
| XP_005090872.1 | HSP70 | *Aplysia californica* |
| XP_005098012.1 | HSP70 | *Aplysia californica* |
| AAB95297.1 | HSP70 | *Biomphalaria glabrata* |
| AAB99911.1 | HSP70 | *Biomphalaria glabrata* |
| NP_001298215.1 | HSP70 | *Biomphalaria glabrata* |
| XP_013072147.1 | HSP70 | *Biomphalaria glabrata* |
| XP_013096386.1 | HSP70 | *Biomphalaria glabrata* |
| XP_013096384.1 | HSP70 | *Biomphalaria glabrata* |
| XP_013096383.1 | HSP70 | *Biomphalaria glabrata* |
| XP_013082115.1 | HSP70 | *Biomphalaria glabrata* |
| XP_013082114.1 | HSP70 | *Biomphalaria glabrata* |
| XP_013081556.1 | HSP70 | *Biomphalaria glabrata* |
| XP_013068425.1 | HSP70 | *Biomphalaria glabrata* |
| XP_013068424.1 | HSP70 | *Biomphalaria glabrata* |
| XP_013091813.1 | HSP70 | *Biomphalaria glabrata* |
| XP_013088343.1 | HSP70 | *Biomphalaria glabrata* |
| XP_013088334.1 | HSP70 | *Biomphalaria glabrata* |
| XP_013096370.1 | HSP70 | *Biomphalaria glabrata* |
| XP_013095344.1 | HSP70 | *Biomphalaria glabrata* |
| XP_013063534.1 | HSP70 | *Biomphalaria glabrata* |
| ALV23100.1 | HSP70 | *Mycoplasma sp.* |
| ABC54952.1 | HSP70 | *Haliotis discus hannai* |
| XP_009056438.1 | HSP70 | *Lottia gigantea* |
| XP_009056460.1 | HSP70 | *Lottia gigantea* |
| XP_009052484.1 | HSP70 | *Lottia gigantea* |
| XP_009051718.1 | HSP70 | *Lottia gigantea* |
| XP_009051716.1 | HSP70 | *Lottia gigantea* |
| XP_009046364.1 | HSP70 | *Lottia gigantea* |
| XP_009045593.1 | HSP70 | *Lottia gigantea* |
| XP_009056443.1 | HSP70 | *Lottia gigantea* |
| AGY78334.1 | HSP70 | *Pomacea canaliculata* |
| AIZ03406.1 | HSP70 | *Pomacea canaliculata* |
| AYH91695.1 | HSP70 | *Pomacea canaliculata* |
| XP_025100818.1 | HSP70 | *Pomacea canaliculata* |
| XP_025100805.1 | HSP70 | *Pomacea canaliculata* |
| XP_025099490.1 | HSP70 | *Pomacea canaliculata* |
| XP_025093615.1 | HSP70 | *Pomacea canaliculata* |
| XP_005113197.2 | HSP40 (Type-III) | *Aplysia californica* |
| XP_012945225.1 | HSP40 (Type-III) | *Aplysia californica* |
| XP_012937761.1 | HSP40 (Type-III) | *Aplysia californica* |
| XP_005096253.1 | HSP40 (Type-III) | *Aplysia californica* |
| XP_005094332.1 | HSP40 (Type-III) | *Aplysia californica* |
| XP_005094324.1 | HSP40 (Type-III) | *Aplysia californica* |
| XP_005106408.1 | HSP40 (Type-III) | *Aplysia californica* |
| XP_012946258.1 | HSP40 (Type-III) | *Aplysia californica* |
| XP_012935695.1 | HSP40 (Type-III) | *Aplysia californica* |
| XP_005104922.1 | HSP40 (Type-III) | *Aplysia californica* |
| XP_035824628.1 | HSP40 (Type-III) | *Aplysia californica* |
| XP_035824627.1 | HSP40 (Type-III) | *Aplysia californica* |
| XP_013068307.1 | HSP40 (Type-III) | *Biomphalaria glabrata* |
| XP_013094320.1 | HSP40 (Type-III) | *Biomphalaria glabrata* |
| XP_013083446.1 | HSP40 (Type-III) | *Biomphalaria glabrata* |
| XP_013083445.1 | HSP40 (Type-III) | *Biomphalaria glabrata* |
| XP_013083444.1 | HSP40 (Type-III) | *Biomphalaria glabrata* |
| XP_013083443.1 | HSP40 (Type-III) | *Biomphalaria glabrata* |
| XP_013068583.1 | HSP40 (Type-III) | *Biomphalaria glabrata* |
| XP_013088548.1 | HSP40 (Type-III) | *Biomphalaria glabrata* |
| XP_013080882.1 | HSP40 (Type-III) | *Biomphalaria glabrata* |
| XP_013077422.1 | HSP40 (Type-III) | *Biomphalaria glabrata* |
| XP_013066765.1 | HSP40 (Type-III) | *Biomphalaria glabrata* |
| XP_013066893.1 | HSP40 (Type-III) | *Biomphalaria glabrata* |
| XP_009056052.1 | HSP40 (Type-III) | *Lottia gigantea* |
| XP_009065217.1 | HSP40 (Type-III) | *Lottia gigantea* |
| XP_009063139.1 | HSP40 (Type-III) | *Lottia gigantea* |
| XP_009062450.1 | HSP40 (Type-III) | *Lottia gigantea* |
| XP_009062448.1 | HSP40 (Type-III) | *Lottia gigantea* |
| XP_009061292.1 | HSP40 (Type-III) | *Lottia gigantea* |
| XP_009061277.1 | HSP40 (Type-III) | *Lottia gigantea* |
| XP_009060482.1 | HSP40 (Type-III) | *Lottia gigantea* |
| XP_009059106.1 | HSP40 (Type-III) | *Lottia gigantea* |
| XP_009058598.1 | HSP40 (Type-I) | *Lottia gigantea* |
| XP_009051357.1 | HSP40 (Type-III) | *Lottia gigantea* |
| XP_009049738.1 | HSP40 (Type-III) | *Lottia gigantea* |
| XP_009049131.1 | HSP40 (Type-III) | *Lottia gigantea* |
| PVD33075.1 | HSP40 (Type-III) | *Pomacea canaliculata* |
| PVD33770.1 | HSP40 (Type-III) | *Pomacea canaliculata* |
| PVD29572.1 | HSP40 (Type-III) | *Pomacea canaliculata* |
| PVD27093.1 | HSP40 (Type-III) | *Pomacea canaliculata* |
| PVD25622.1 | HSP40 (Type-III) | *Pomacea canaliculata* |
| PVD24366.1 | HSP40 (Type-III) | *Pomacea canaliculata* |
| PVD20593.1 | HSP40 (Type-II) | *Pomacea canaliculata* |
| XP_005108410.1 | HSP20 | *Aplysia californica* |
| XP_005108414.1 | HSP20 | *Aplysia californica* |
| XP_013090183.1 | HSP20 | *Biomphalaria glabrata* |
| XP_013085554.1 | HSP20 | *Biomphalaria glabrata* |
| XP_013076884.1 | HSP20 | *Biomphalaria glabrata* |
| XP_013069954.1 | HSP20 | *Biomphalaria glabrata* |
| XP_013067909.1 | HSP20 | *Biomphalaria glabrata* |
| XP_013073434.1 | HSP20 | *Biomphalaria glabrata* |
| XP_013092927.1 | HSP20 | *Biomphalaria glabrata* |
| AMX23358.1 | HSP20 | *Haliotis discus hannai* |
| ABR57318.1 | HSP20 | *Haliotis discus hannai* |
| XP_009066881.1 | HSP20 | *Lottia gigantea* |
| XP_009056339.1 | HSP20 | *Lottia gigantea* |
| XP_009056338.1 | HSP20 | *Lottia gigantea* |
| XP_009055472.1 | HSP20 | *Lottia gigantea* |
| XP_009045112.1 | HSP20 | *Lottia gigantea* |
| AYH91698.1 | HSP20 | *Pomacea canaliculata* |
| XP_005097388.1 | HSP10 | *Aplysia californica* |
| XP_013062221.1 | HSP10 | *Biomphalaria glabrata* |
| XP_009045000.1 | HSP10 | *Lottia gigantea* |
| XP_025095803.1 | HSP10 | *Pomacea canaliculata* |

Table S2 List of Genbank accession numbers and download links of RNA-Seq data in five tissues of *P. canaliculata*

| GenBank Accession | Size(Mb) | Tissues | Download Links |
| --- | --- | --- | --- |
| SRR6429145 | 3257 | Hepatopancreas | https://sra-downloadb.be-md.ncbi.nlm.nih.gov/sos1/sra-pub-run-2/SRR6429145/SRR6429145.1 |
| SRR6429146 | 3329 | Hepatopancreas | https://sra-downloadb.be-md.ncbi.nlm.nih.gov/sos1/sra-pub-run-2/SRR6429146/SRR6429146.1 |
| SRR6429153 | 1025 | Hepatopancreas | https://sra-downloadb.be-md.ncbi.nlm.nih.gov/sos2/sra-pub-run-13/SRR6429153/SRR6429153.1 |
| SRR6429132 | 3380 | Kidney | https://sra-downloadb.be-md.ncbi.nlm.nih.gov/sos1/sra-pub-run-2/SRR6429132/SRR6429132.1 |
| SRR6429133 | 3409 | Kidney | https://sra-downloadb.be-md.ncbi.nlm.nih.gov/sos2/sra-pub-run-13/SRR6429133/SRR6429133.1 |
| SRR6429162 | 1000 | Kidney | https://sra-downloadb.be-md.ncbi.nlm.nih.gov/sos1/sra-pub-run-2/SRR6429162/SRR6429162.1 |
| SRR6429154 | 3261 | Gill | https://sra-downloadb.be-md.ncbi.nlm.nih.gov/sos1/sra-pub-run-2/SRR6429154/SRR6429154.1 |
| SRR6429140 | 3332 | Gill | https://sra-downloadb.be-md.ncbi.nlm.nih.gov/sos1/sra-pub-run-2/SRR6429140/SRR6429140.1 |
| SRR6429141 | 766 | Gill | https://sra-downloadb.be-md.ncbi.nlm.nih.gov/sos2/sra-pub-run-13/SRR6429141/SRR6429141.1 |
| SRR6429161 | 3252 | Ovary | https://sra-downloadb.be-md.ncbi.nlm.nih.gov/sos2/sra-pub-run-13/SRR6429161/SRR6429161.1 |
| SRR6429160 | 3329 | Ovary | https://sra-downloadb.be-md.ncbi.nlm.nih.gov/sos2/sra-pub-run-13/SRR6429160/SRR6429160.1 |
| SRR6429159 | 1213 | Ovary | https://sra-downloadb.be-md.ncbi.nlm.nih.gov/sos2/sra-pub-run-13/SRR6429159/SRR6429159.1 |
| SRR6429142 | 3293 | Testis | https://sra-downloadb.be-md.ncbi.nlm.nih.gov/sos1/sra-pub-run-2/SRR6429142/SRR6429142.1 |
| SRR6429143 | 3374 | Testis | https://sra-downloadb.be-md.ncbi.nlm.nih.gov/sos2/sra-pub-run-13/SRR6429143/SRR6429143.1 |
| SRR6429144 | 726 | Testis | https://sra-downloadb.be-md.ncbi.nlm.nih.gov/sos2/sra-pub-run-13/SRR6429144/SRR6429144.1 |

Table S3 Primers of the PcaHSP genes for qPCR

| Gene | Primer Name | Primer Sequence（5’→3’） | Annealing temperature in qPCR |
| --- | --- | --- | --- |
| *PcaHSP90-1* | PcaHsp90-1-F | ATGTTACTATGCAGCATGGTG | 60℃ |
|  | PcaHsp90-1-R | TAGTCTGTGTCACTAAGGAAG |  |
| *PcaHSP90-2* | PcaHsp90-2-F | ATCTGGAACATATGAGGTGAG | 60℃ |
|  | PcaHsp90-2-R | GTAGGATATTGACACGATGAC |  |
| *PcaHSP70-1* | PcaHsp70-1-F | AACTTCTGGACATGCATGACTG | 60℃ |
|  | PcaHsp70-1-R | ACACTTTTGTGGTGAAGCTTGAG |  |
| *PcaHSP70-2* | PcaHsp70-2-F | AGCTTTCTTCACGGTTAGCATC | 60℃ |
|  | PcaHsp70-2-R | ACGTCAAGCAGATTTTGGCTAAC |  |
| *PcaHSP70-3* | PcaHsp70-3-F | ACAACACCCATCATGTGATCCA | 60℃ |
|  | PcaHsp70-3-R | TGGCCTATGGTCTACACAAGAAAG |  |
| *PcaHSP70-4* | PcaHsp70-4-F | TGGGTACAACATACTCCTGTGTTG | 60℃ |
|  | PcaHsp70-4-R | TGCTTCATGTCTGATTGGAC |  |
| *PcaHSP70-5* | PcaHsp70-5-F | TCTTCCTTTTCACCATTTACC | 53℃ |
|  | PcaHsp70-5-R | TGCAACGTGTTATTTCAGAC |  |
| *PcaHSP70-6* | PcaHSP70-6-F | TGTCTTTCTTGTACTTGCGTTG | 60℃ |
|  | PcaHSP70-6-R | AATGTCCTCATCTTTGACTTG |  |
| *PcaHSP70-7* | PcaHsp70-7-F | AAGGCAAAGGAGAGAAGAATG | 60℃ |
|  | PcaHsp70-7-R | TTCTTGTACTTGCGTTGGAAC |  |
| *PcaHSP70-8* | PcaHsp70-8-F | TCACCTCGATCTGAGGAACAC | 60℃ |
|  | PcaHsp70-8-R | ATCGCAGATCTTCACCACC |  |
| *PcaHSP70-9* | PcaHsp70-9-F | AGTGCCATGGAGCTTAGTC | 60℃ |
|  | PcaHsp70-9-R | TGACAAACTGAGTAGCGCC |  |
| *PcaHSP70-10* | PcaHsp70-10-F | TCTTCAGTTTGCTGTACGAC | 60℃ |
|  | PcaHsp70-10-R | AACCTCTGTGCCTTGTTTTC |  |
| *PcaHSP70-11* | PcaHsp70-11-F | ACGTACAGACAAGAAGTAC | 60℃ |
|  | PcaHsp70-11-R | CTTCATGTTGAATACGTCTC |  |
| *PcaHSP70-12* | PcaHsp70-12-F | TAGCATTCAGGAATGGAGAAC | 60℃ |
|  | PcaHsp70-12-R | TTGAACAGTACAGTTCCTCTC |  |
| *PcaHSP70-13* | PcaHsp70-13-F | ACATAGCTGGGAGTAGTTCTC | 60℃ |
|  | PcaHsp70-13-R | GGATACTGTGAAACGGAATG |  |
| *PcaHSP60-1* | PcaHsp60-1-F | TCACTAGCTAGATGCTATG | 60℃ |
|  | PcaHsp60-1-R | CTTTTGTGATCTTCGGACTTC |  |
| *PcaHSP40-1* | PcaHsp40-1-F | TGATCTCGTCTGATGATG | 60℃ |
|  | PcaHsp40-1-R | TTCTCGCCAATGTTGACTC |  |
| *PcaHSP40-2* | PcaHsp40-2-F | AGTTAAGGTATCTGTTCGCCAAG | 60℃ |
|  | PcaHsp40-2-R | TGAATCCCTTGTAACTCCAAG |  |
| *PcaHSP40-3* | PcaHsp40-3-F | TACTGTCCCATCAACTAAG | 60℃ |
|  | PcaHsp40-3-R | CAGTCACATGATCGAATTC |  |
| *PcaHSP40-4* | PcaHsp40-4-F | TCATAAGAGCAGGTTCTGTG | 60℃ |
|  | PcaHsp40-4-R | TGGCAAAAGCTAAAGAGGTC |  |
| *PcaHSP40-7* | PcaHsp40-7-F  PcaHsp40-7-R | ATGATCTTACTTTGGCTGCTC  TCTCAAATTCCTTGTCTCGTG | 60℃ |
| *PcaHSP40-8* | PcaHsp40-8-F | TCCTCATAATCCACTAGATTC | 60℃ |
|  | PcaHsp40-8-R | AGGCATGCCAATATACAGAAAC |  |
| *PcaHSP40-9* | PcaHsp40-9-F | TGGGACTTCAAGAATTTCATC | 60℃ |
|  | PcaHsp40-9-R | GTATTACGGACCTGAAAATAC |  |
| *PcaHSP40-10* | PcaHsp40-10-F | TGAAGCTGTTTTGATCGCATG | 60℃ |
|  | PcaHsp40-10-R | TTCAGCAACTTTTGGCACTG |  |
| *PcaHSP40-11* | PcaHsp40-11-F | ACAGATTGATGTGAAACCTG | 60℃ |
|  | PcaHsp40-11-R | CACAAAGTGCATCACGTAAAG |  |
| *PcaHSP40-12* | PcaHsp40-12-F | TCTACATTTTGATCTGGCAG | 60℃ |
|  | PcaHsp40-12-R | AAGGCTCACCACTATAGAAAC |  |
| *PcaHSP40-13* | PcaHsp40-13-F | TTCCTTTTCGTCCAGTACTATG | 60℃ |
|  | PcaHsp40-13-R | TCTGCAATGGCCAGAAAATAG |  |
| *PcaHSP20-1* | PcaHsp20-1-F | AGTCCTTTCCGAAGTCTTTTC | 60℃ |
|  | PcaHsp20-1-R | TTCAGCAGCATGTTCAAGGATG |  |
| *PcaHSP20-2* | PcaHsp20-2-F | AAGAGTTCTTCGTGCTCAAG | 60℃ |
|  | PcaHsp20-2-R | TGACGATGAGCTTGTTCTC |  |
| *PcaHSP20-3* | PcaHsp20-3-F | ATCAGAGACACGTTCAATC | 60℃ |
|  | PcaHsp20-3-R | CAACTTTGACCTTCAGTTC |  |
| *PcaHSP20-4* | PcaHsp20-4-F | TGTGCGCATTGATGTCAAGC | 62℃ |
|  | PcaHsp20-4-R | ACCGCACTCTTGTCAACATG |  |
| *PcaHSP20-5* | PcaHsp20-5-F | AACCTGAAGAGATCAACATC | 60℃ |
|  | PcaHsp20-5-R | AACACTCAGAGATGATGTG |  |
| *PcaHSP20-6* | PcaHsp20-6-F | ACGTACTGACGAGTGAATTGTC | 60℃ |
|  | PcaHsp20-6-R | ATGATGGTACCCAGAATG |  |
| *PcaHSP20-7* | PcaHsp20-7-F | ACCGTCCTTTCACTAGTATC | 60℃ |
|  | PcaHsp20-7-R | TGCTTAGCATGGATGATGAG |  |
| *PcaHSP20-8* | PcaHsp20-8-F | AGACTTCTTTAATCCACAGAG | 60℃ |
|  | PcaHsp20-8-R | ACAGGCATTGATGATAACAC |  |
| *PcaHSP20-9* | PcaHsp20-9-F | AGGTCTTCTGGTATGTAATG | 60℃ |
|  | PcaHsp20-9-R | TTCATCATGAGAGAGTTCAC |  |
| *PcaHSP20-10* | PcaHsp20-10-F | AAGCCAGTGAACACTTGTC | 60℃ |
|  | PcaHsp20-10-R | AGCGAAAGACACAAAGTTTGAG |  |
| *PcaHSP20-11* | PcaHsp20-11-F | TTTTGCAGAACTCTGGTGCA | 60℃ |
|  | PcaHsp20-11-R | AAGGGGAGGAGAAAGCTTGAAAA |  |
| *PcaHSP10-1* | PcaHsp10-1-F | TGGTTTCTGTCACAACTC | 60℃ |
|  | PcaHsp10-1-R | GTCCAAGTTGGTGATAAG |  |
| *β-actin* | qactin-F | TCACCATTGGCAACGAGCGAT |  |
|  | qactin-R | TCTCGTGAATACCAGCCGACT |  |

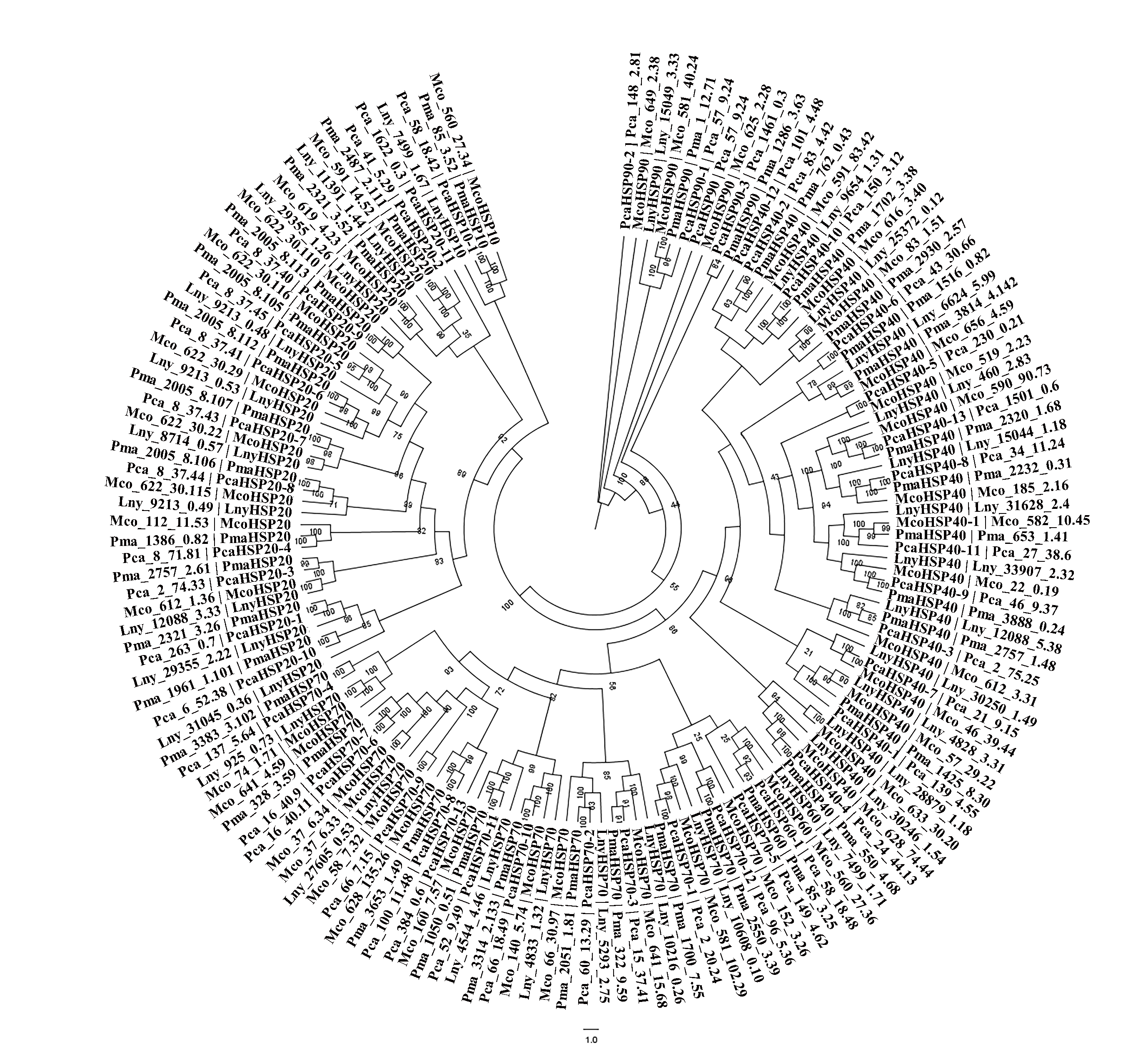

Fig. S1. Phylogenetic relationships of HSP proteins from *P. canaliculata*, *P. maculata*, *L. nyassanus*, and *M. cornuarietis*.

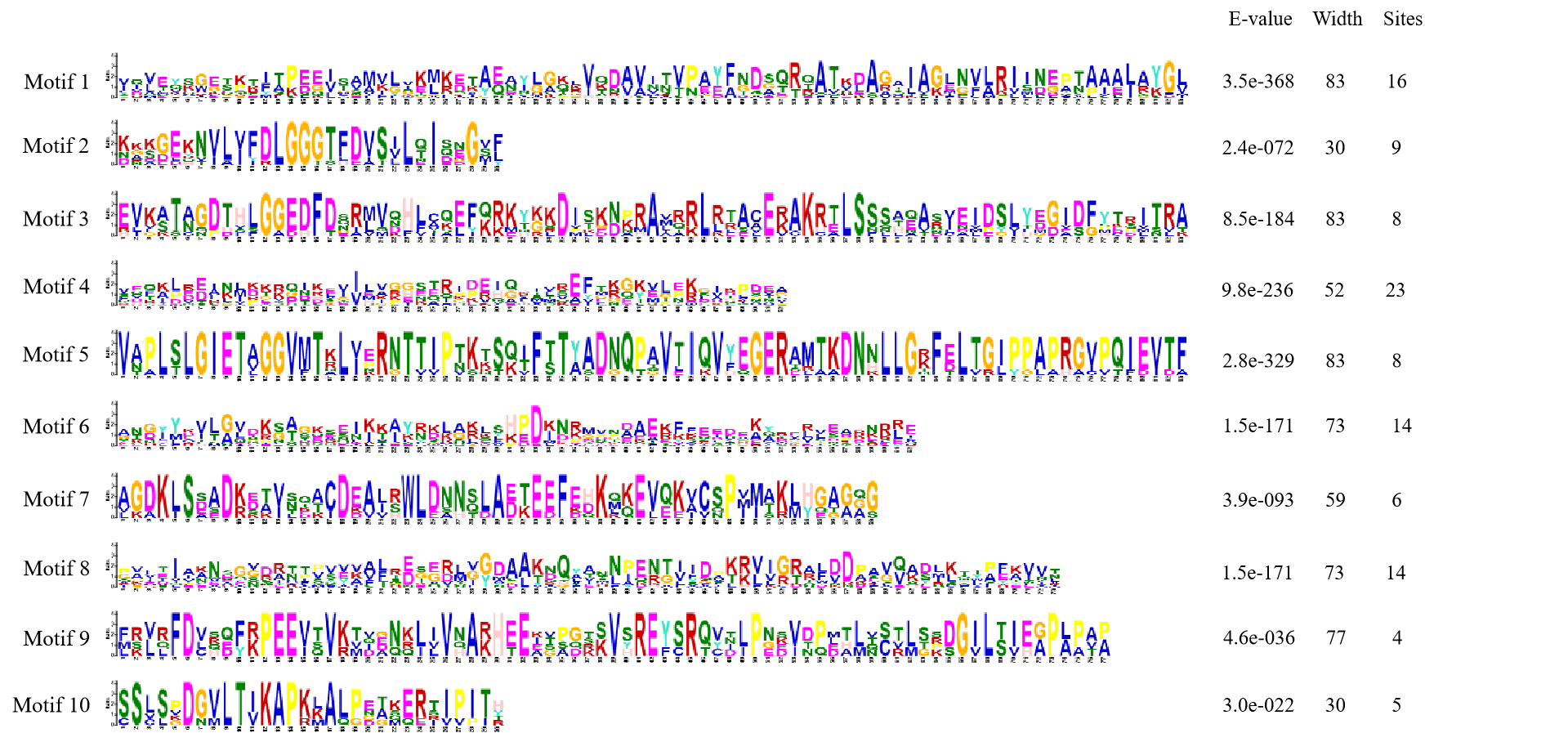

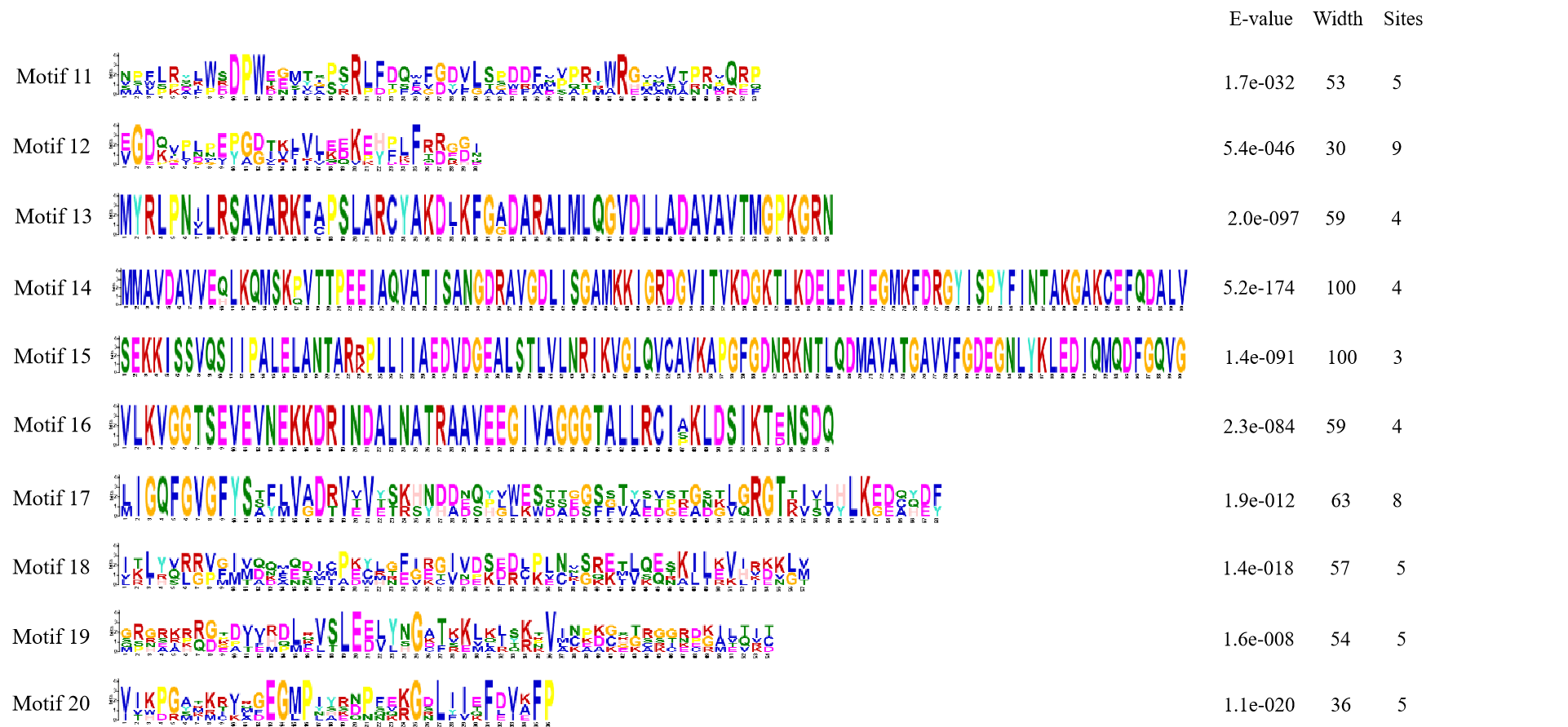

Fig. S2. Sequence logos for the conserved motifs of PcaHSP proteins.

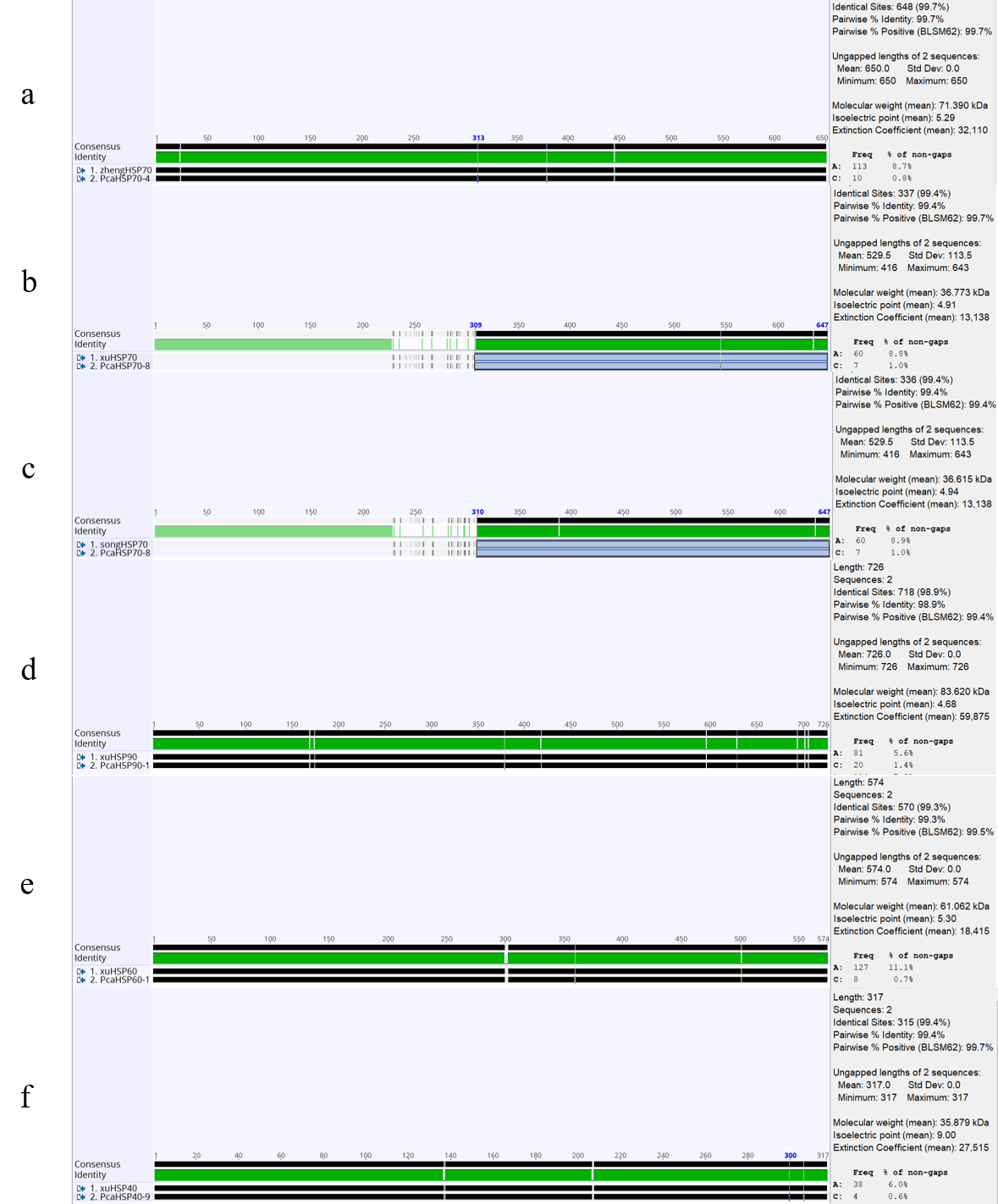

Fig. S3. The results of published Hsp protein sequences of *P. canaliculata* compared with the PcaHSP families identified in this study using Geneious 11.1.5. (A) Zheng et al. , (2012) HSP70 was 99.7% similarity with PcaHSP70-4. (B) Xu et al. (2014) HSP70 was 99.4% similarity with PcaHSP70-8. (C) Song et al. (2014) HSP70 was 99.4% similarity with PcaHSP70-8. (D) Xu et al. (2014) HSP90 was 97.6% similarity with PcaHSP90-1 . (E) Xu et al. (2014) HSP60 was 99.5% similarity with PcaHSP60-1. (F) Xu et al. (2019) HSP40 was 98% similarity with PcaHSP40-9.
